# Admixture of evolutionary rates across a hybrid zone

**DOI:** 10.1101/2021.09.27.461223

**Authors:** Tianzhu Xiong, Xueyan Li, Masaya Yago, James Mallet

## Abstract

Hybridization is a major evolutionary force that can erode genetic differentiation between species, whereas reproductive isolation maintains such differentiation. In studying a hybrid zone between the swallowtail butterflies *Papilio syfanius* and *Papilio maackii*, we made the unexpected discovery that genomic substitution rates are unequal between the parental species. This phenomenon creates a novel process in hybridization, where genomic regions most affected by gene flow evolve at similar rates, while genomic regions with greater reproductive isolation evolve at divergent rates. Thus, hybridization mixes evolutionary rates in a way similar to its effect on ancestry. Using coalescent theory, we show that the rate-mixing process provides distinct information about levels of gene flow across different parts of genomes, and that maintenance of divergent substitution rates can be predicted quantitatively from relative sequence divergence (*F*_*ST*_) between the hybridizing species at equilibrium. A corollary is that divergent rates will be maintained in regions linked to barrier loci. Overall, we demonstrate that reproductive isolation maintains not only the final outcome of genomic differentiation, but also the rate at which differentiation accumulates. This new information also suggests that the separation of evolutionary rates co-localizes with the separation of gene pools between genomes of incipient species.

## 1. Introduction

DNA substitution, where single-nucleotide mutations accumulate through time, is a critical process in molecular evolution. Both molecular phylogenetics and coalescent theory rely on observed mutations to reconstruct gene genealogies (Wakeley, 2016; Yang and Rannala, 2012), and so the rate of substitution/mutation is the predominant link from molecular data to information about the timing of past events (Bromham and Penny, 2003). Substitution rates often vary among lineages: generation time, spontaneous mutation rate, and fixation probabilities of new mutations could all contribute to the variation of substitution rates (Lynch, 2010; Ohta, 1993). Recent evidence even suggests mutation rates are variable among human populations (DeWitt et al., 2021; Harris, 2015). As such variation affects how fast the molecular clock ticks, reconstructing gene genealogies among different species usually accounts for species-specific rates of evolution (Lepage et al., 2007). However, under the standard coalescent framework, empirical studies of within- and between-species variation tend to ignore rate variation among populations (Costa and Wilkinson-Herbots, 2017; Kautt et al., 2020; Wolf and Ellegren, 2017). The latter is partly based on a popular assumption in coalescent theory that neutral mutation rate is constant for a given site across the whole genealogy (Hudson, 1990). Hybridization and speciation lie in the gray zone of these extremes, and have their own problem: molecular clocks from different lineages could be coupled by cross-species gene flow — a gene could evolve under one clock before it flows into another species and switches to evolve according to a different clock (Fig. 1). This coupling process is largely outside the scope of existing theories, and has received little attention from empirical studies.

**Figure 1:**
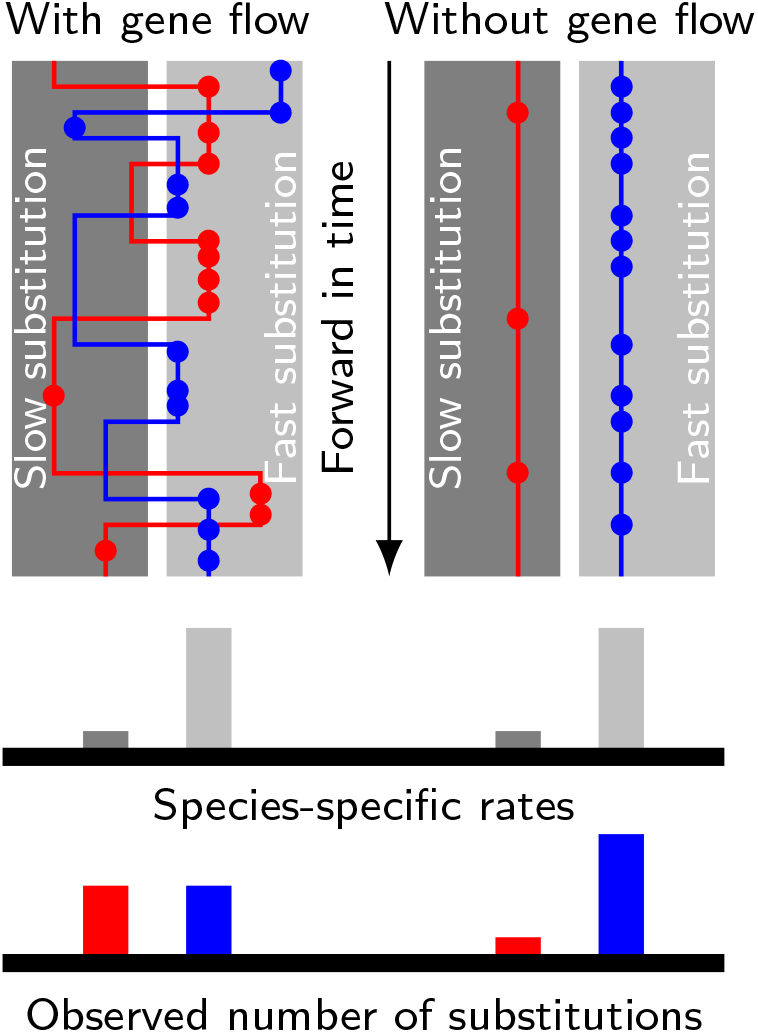
Gene flow interacts with divergent substitution rates and affects observed numbers of substitutions. Each gray block represents a species with its species-specific substitution rate. Solid lines represent gene genealogies prior to coalescence, and horizontal jumps between species represent interspecific gene flow. Dots are substitutions. When a gene sequence inherits mutations derived under multiple rates, the number of substitutions it carries will reflect a mixture of substitution rates among different species. If gene flow is strong, each lineage carries a similar number of substitutions; If gene flow is weak, genes evolve independently with species-specific rates, and the distribution of substitutions in each lineage will likely be skewed towards the distribution of species-specific substitution rates.

However, if coupling between molecular clocks exists and can be measured, its strength could carry information about gene flow, which is important for studying reproductive isolation between incipient species. It has been shown that genes responsible for reproductive isolation lead to locally elevated genomic divergence (“genomic islands”), often caused by linked genomic regions experiencing less gene flow (“barrier loci”) (Michel et al., 2010; Nosil et al., 2009; Payseur and Rieseberg, 2016; Renaut et al., 2013). In studying a hybrid zone between two butterfly species, *Papilio syfanius* and *Papilio maackii*, we discovered evidence for unequal genome-wide substitution rates between the two species. Using this system, we investigate the interaction between unequal substitution rates and gene flow, and whether this interaction reveals new information on reproductive isolation.

As these butterflies are rare species, occurring in a remote region of China, and are hard to collect, we employed methods based on analysis of whole-genome sequences of a few specimens. We hope that these methods may prove of use in studying other rare or perhaps endangered species where few individuals can be sacrificed. Results will follow two parallel lines: first, we provide evidence that genomic islands are associated with barrier loci. Then we infer the existence of unequal substitution rates. Finally, using a coalescent model, we calculate the relationship between the magnitude of genomic islands and the degree of coupling between substitution rates in linked regions. Throughout the analysis, we assume no reverse mutations, so that higher substitution rates always lead to higher numbers of observed substitutions.

## 2. Results

### 2.1 Divergent sister species with ongoing hybridization

We sampled 11 males of *P. syfanius* & *P. maackii* across a geographic transect covering both pure populations (in the sense of being geographically away from the hybrid zone) and hybrid populations (Fig. 2A, dashed lines). We also include four outgroup species, two of which have chromosome-level genome assemblies (*P. bianor*, *P. xuthus*) (Li et al., 2015; Lu et al., 2019), while the other two (*P. arcturus*, *P. dialis*) are new to this study. All samples were re-sequenced to at least 20× coverage across the genome and mapped to the genome assembly of *P. bianor*. Among sampled local populations, *P. syfanius* inhabits the highlands of Southwest China (Fig. 2A, red region), while *P. maackii* dominates at lower elevations (Fig. 2A, blue region). The two lineages form a spatially contiguous hybrid zone at the edge of the Hengduan Mountains (Fig. 2B) with individuals exhibiting intermediate wing patterns (Fig. 2A: purple dot, corresponding to population WN in Fig. 2B). Consistent with previous results (Condamine et al., 2013), assembled whole mitochondrial genomes are not distinct between the two lineages (Fig. 2C), suggesting either that divergence was recent, or that gene flow has homogenized the mitochondrial genomes. However, the two species are likely adapted to different environments associated with altitude, as several ecological traits are strongly divergent (Kashiwabara, 1991) (Fig. S1). Similarly, between pure populations (KM & XY in Fig. 2B), relative divergence (*F*_*ST*_) is also high across the entire nuclear genome (Fig. 2D). The *F*_*ST*_ on autosomes averages between 0.2 − 0.4, and on the sex chromosome (Z-chromosome) it reaches 0.78. A highly heterogeneous landscape of *F*_*ST*_ is accompanied by numerous islands of elevated sequence divergence (*D_XY_*) and reduced genetic diversity (*π*) scattered across the genome (Fig. S2). Overall, despite ongoing hybridization, genomes of the pure populations of *P. syfanius* and *P. maackii* are strongly differentiated.

**Figure 2:**
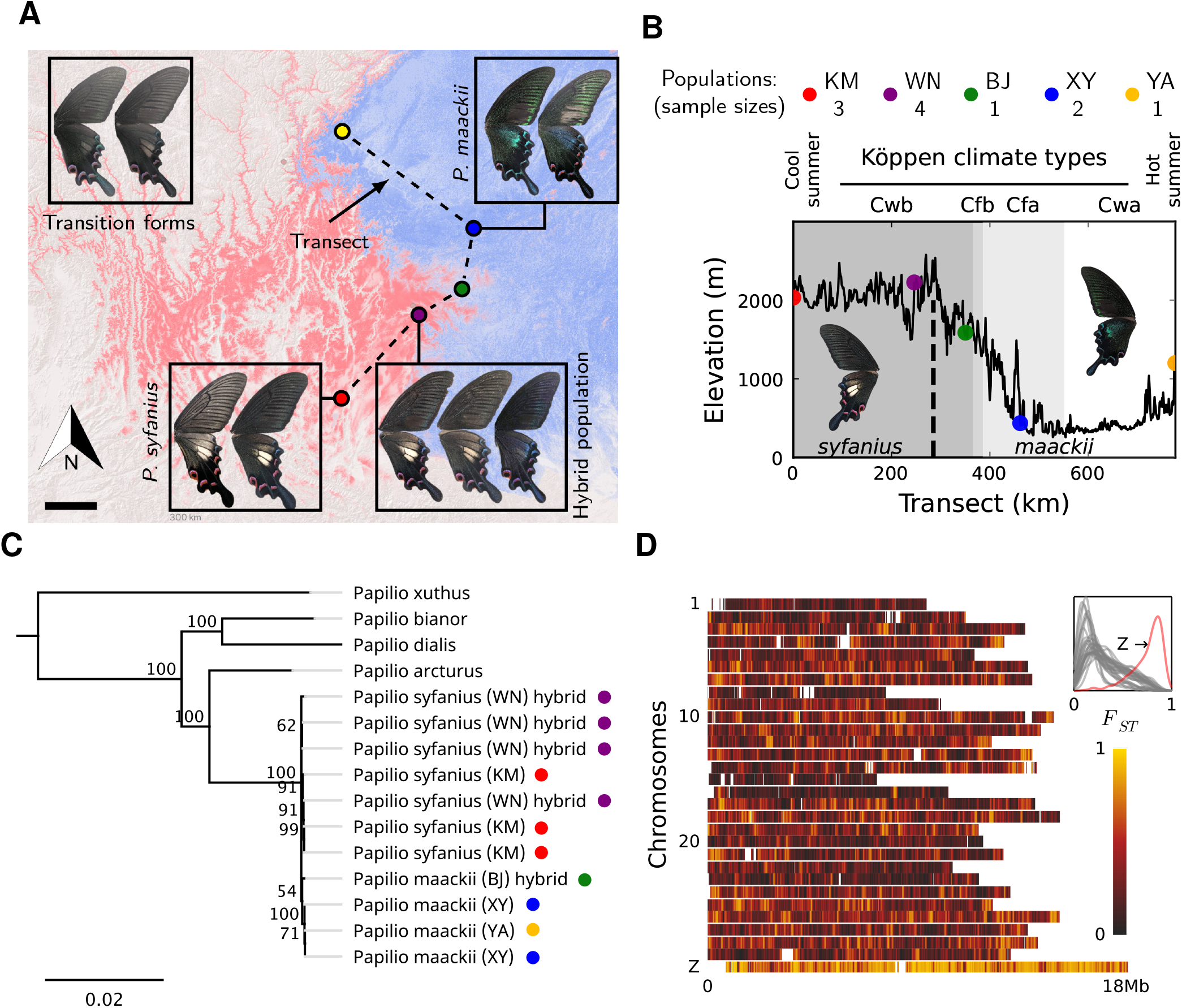
Overview of the study system. **(A)** The geographic distribution of *P. syfanius* (red) and *P. maackii* (blue). Scale-bar: 100km. Dashed line is the sampling transect covering five populations (colored circles). **(B)** Elevation, climate, and sample sizes along the transect. **(C)** Mitochondrial tree with four outgroups. **(D)** *F*_*ST*_ across chromosomes (50kb windows with 10kb increments). The inset shows the estimated density of *F*_*ST*_ on each chromosome.

### 2.2 Genomic islands are associated with barrier loci

A natural question is whether genomic differentiation is associated with barrier loci and reproductive isolation. In other words, can *F*_*ST*_ variation be attributed to gene flow variation between sister species? We suspect that barrier loci likely exist, because sequence variation between pure populations suggests that elevated *F*_*ST*_ is associated with reduced *π* and elevated *D*_*XY*_ across autosomes (Fig. 3A), as expected for hybridizing species (Irwin et al., 2018). The Z chromosome (sex chromosome) has the highest *D*_*XY*_ and the lowest *π* among all chromosomes (Fig. S2), another characteristic of hybridizing species with barriers to gene flow (Kronforst et al., 2013).

**Figure 3:**
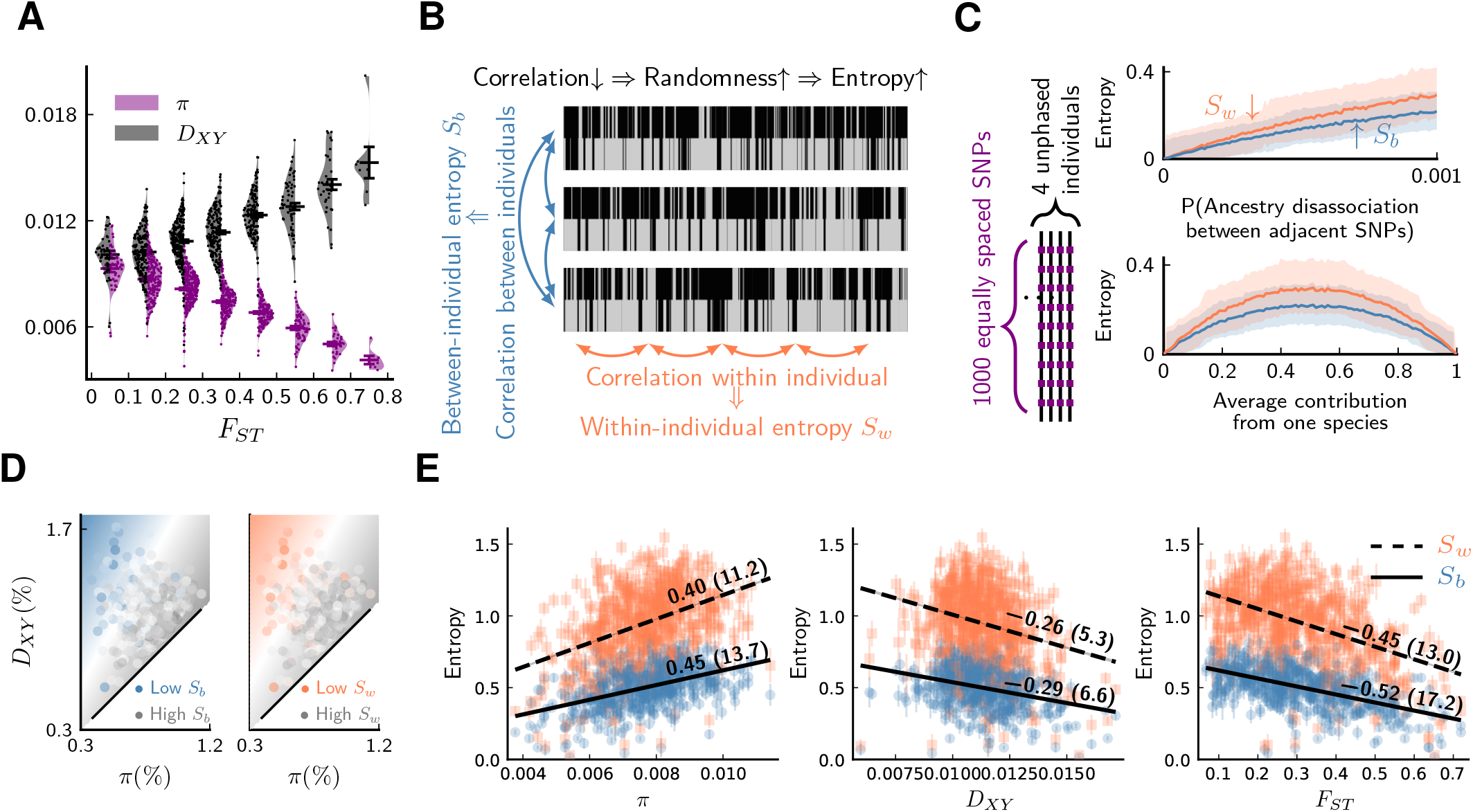
Evidence of barrier loci on autosomes. For all plots, pure populations refer to XY & KM, and hybrid population refers to WN. **(A)** Between pure populations, reduced diversity (*π*, showing values averaged between pure populations) is associated with increased divergence (*D*_*XY*_) across autosomes (30 segments per chromosome). Error bars are standard errors. **(B)** The conceptual picture of entropy metrics on diploid, unphased ancestry signals. In a genomic window, between-individual entropy (*S*_*b*_) measures local ancestry randomness among individuals, while within-individual entropy (*S_w_*) measures ancestry randomness along a chromosomal interval. They carry different information: for example, in a hybrid swarm (Hasselman et al., 2014) subject to long-term inbreeding, *S*_*b*_ will drop to zero because ancestry completely correlates between individuals, but *S_w_* could be large due to recombination breaking up ancestry tracts. **(C)** Simulated behaviors of entropy in a simplified model of biparental local ancestry. Chromosomes are assumed to be spatially homogeneous, thus recombination rate is uniform among 1000 equally spaced SNPs, and adjacent SNPs have a single probability of ancestry disassociation. For each haplotype block with linked ancestry, its ancestry is randomly assigned according to the average contribution from each species. Each pair of haploid chromosomes were combined into an unphased ancestry signal before calculating entropy. The top plot assumes equal contribution from both species, and the bottom plot assumes ancestry disassociation probability = 0.001. Solid lines are average entropies across 1000 repeated simulations, and shaded areas represent upper and lower average deviations from the mean. **(D)** The joint range among entropy, *π*, and *D*_*XY*_ across autosomes (20 segments per chromosome). Color range is normalized by the range of entropy in each plot. Gray represents higher entropy, and colored regions are associated with lower entropy. Heatmaps represent linear fits to the ensembles of points. **(E)** The correlation *ρ* on autosomes between entropy in hybrid populations and {*π*, *D*_*XY*_, *F*_*ST*_} within and between pure populations. *ρ* is shown above each regression line. Error bars are standard errors of entropy from 50 repeated estimates of local ancestry using software ELAI (parameters are in Materials and Methods). The significance of *ρ* was estimated using block-jackknifing among all segments: *Z*-scores are shown in parentheses.

To strengthen evidence for barrier loci, we augment the analysis with the sequences of four individuals from the population closest to the center of the hybrid zone (Population WN). We investigate whether differences in ancestry variation provide additional evidence for barrier loci in this hybrid zone. The underlying logic is that barrier loci will simultaneously:

1. Reduce linked *π* in pure populations (Ravinet et al., 2017);
2. Elevate linked *D*_*XY*_ between pure populations (Ravinet et al., 2017);
3. Elevate linkage disequilibrium in hybrid zones (Barton, 1983);
4. Enrich linked ancestry from one lineage in hybrid zones (Sedghifar et al., 2016).

Effects 3 and 4 can be bundled together as “reduced ancestry randomness” around barrier loci because both are expected if intermixing of segments of different ancestries within a genomic interval is prevented. For effects 3 and 4, because of small sample sizes, estimating site-specific statistics such as pairwise linkage disequilibrium is untenable. However, our high-quality chromosome-level reference genome enabled accurate estimation of local ancestry. As a remedy for small sample sizes, we employ two entropy metrics borrowed from signal-processing theory to quantify ancestry randomness in local regions along chromosomes (see Materials and Methods). By dividing chromosomes into segments, we can extract indirect information about effects 3 and 4 at the expense of reduced genomic resolution. The proposed metrics, *S*_*b*_ and *S*_*w*_, correspond to the randomness of ancestry between and within individual diploid genomes from a local population (Fig. 3B). For a cohort of ideal chromosomes with uniform recombination and marker density, if ancestry is independent between homologous chromosomes, both *S*_*b*_ and *S*_*w*_ increase with reduced local ancestry correlation and more balanced parental contribution (Fig. 3C).

For a given autosomal segment, we then calculate *π*, *D*_*XY*_, and *F*_*ST*_ between pure populations, as well as entropy metrics *S*_*b*_ and *S*_*w*_ in hybrid individuals for the same segment. To investigate whether effects 1-4 are all present in our system, the joint range among entropy, *π*, and *D*_*XY*_ is shown in Fig. 3D, which suggests that low ancestry randomness (low entropy) is likely associated with reduced *π* within species and elevated *D*_*XY*_ between species. To further quantify such association, we estimated Pearson’s correlation coefficients (*ρ*) between entropy and the latter statistics (Fig. 3E). These associations are strongly significant (*Z*-scores ≫ 3). Consequently, reduced ancestry randomness in hybrids (effects 3 & 4) coincides with classical patterns of barrier loci between pure populations (effects 1 & 2). This analysis is not sufficient to exclude all alternative hypotheses. For instance, we cannot entirely exclude the possibility that patterns are driven by low-recombination regions (*S*_*w*_, *S*_*b*_ ↓) experiencing linked selection (*π* ↓) and also having elevated mutation rates (*D*_*XY*_ ↑). Nonetheless, this alternative seems most unlikely, as low-recombination regions would often be less rather than more mutable (Arbeithuber et al., 2015; Jensen-Seaman et al., 2004; Lercher and Hurst, 2002; Liu et al., 2016; Yang et al., 2015). Overall, adding information from hybrid populations strengthens the evidence for barrier loci acting across autosomes.

The Z chromosome was excluded from the analysis as it likely differs in mutation rate or effective population size (Presgraves, 2018), but its ancestry in hybrid individuals either retains purity or resembles very recent hybridization (long blocks of heterozygous ancestry, Fig. S5). The Z chromosome has low ancestry randomness, and it also has the highest level of divergence (Fig. 2D), both of which suggest strong barriers to gene flow between *P. syfanius* and *P. maackii* on this chromosome.

### 2.3 Asymmetric site patterns

The first hint of divergent substitution rates between the two species comes from site-pattern asymmetry, although this alone is insufficient to establish the existence of divergent rates. We focus on two kinds of biallelic site patterns. In the first kind, choose three taxa (P_1_,P_2_,O_1_), with P_1_=*syfanius*, P_2_=*maackii*, while O_1_ is an outgroup. Assuming no other factors, if substitution rates are equal between P_1_ and P_2_, then site pattern (P_1_,P_2_,O_1_)=(A,B,B) and (B,A,B) occur with equal frequencies, where “A” and “B” represent distinct alleles. This leads to a *D* statistic describing the asymmetry between three-taxon site patterns:

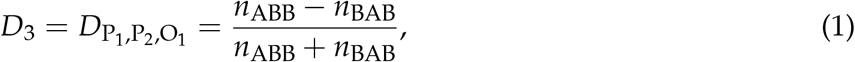

where *n* is the count of a particular site pattern across designated genomic regions. A significantly non-zero *D*_3_ indicates strongly asymmetric distributions of alleles between P_1_ and P_2_.

In a second kind of site pattern test, we compare four taxa (P_1_,P_2_,O_1_,O_2_) and calculate a similar statistic between site patterns (A,B,B,A) vs (B,A,B,A):

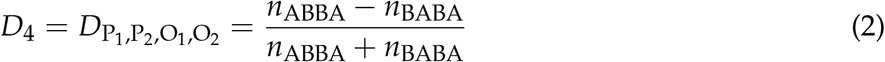

Classically, a significantly non-zero *D*_4_ suggests that gene flow occurs between an outgroup and either P_1_ or P_2_, thus it is widely used to detect hybridization (the ABBA-BABA test) (Durand et al., 2011; Hibbins and Hahn, 2022). Here, *D*_4_ is used more generally as an additional metric of site pattern asymmetry.

We compute both *D*_3_ and *D*_4_ on synonymous, nonsynonymous, and intronic sites, with *P. syfanius* and *P. maackii* samples taken from pure populations (KM & XY). For each type of site, we progressively exclude regions with local *F*_*ST*_ below a certain threshold, and report *D* statistics on the remaining sites in order to show the increasing site-pattern asymmetry in more divergent regions (Fig. 4). *D*_3_ is significantly negative regardless of outgroup or site type for most *F*_*ST*_ thresholds, and *D*_4_ is also significantly negative for most outgroup combinations when computed across the entire genome (*Z*-scores are shown in Fig. S3), proving that site-patterns are strongly asymmetric between *P. syfanius* and *P. maackii*. Importantly, the direction of asymmetry is nearly identical across all outgroup comparisons. This asymmetry cannot be attributed to batch-specific variation as all samples were processed and sequenced in a single run, and variants were always called on all individuals of *P. syfanius* and *P. maackii*. Sequencing coverage is normal for most annotated genes used in the analysis (Table S2), suggesting that asymmetry is not due to systematic copy-number variation that could affect variant calls. Nonetheless, two independent processes of evolution could explain observed asymmetric site patterns. In hypothesis I (Fig. 5A, left), asymmetry is generated via stronger gene flow between *P. syfanius* and outgroups, leading to biased allele-sharing. In hypothesis II, site pattern asymmetry is due to unequal substitution rates between *P. syfanius* and *P. maackii*, which is further modified by recurrent mutations in all four outgroups (Fig. 5A, right). We test each hypothesis below.

**Figure 4:**
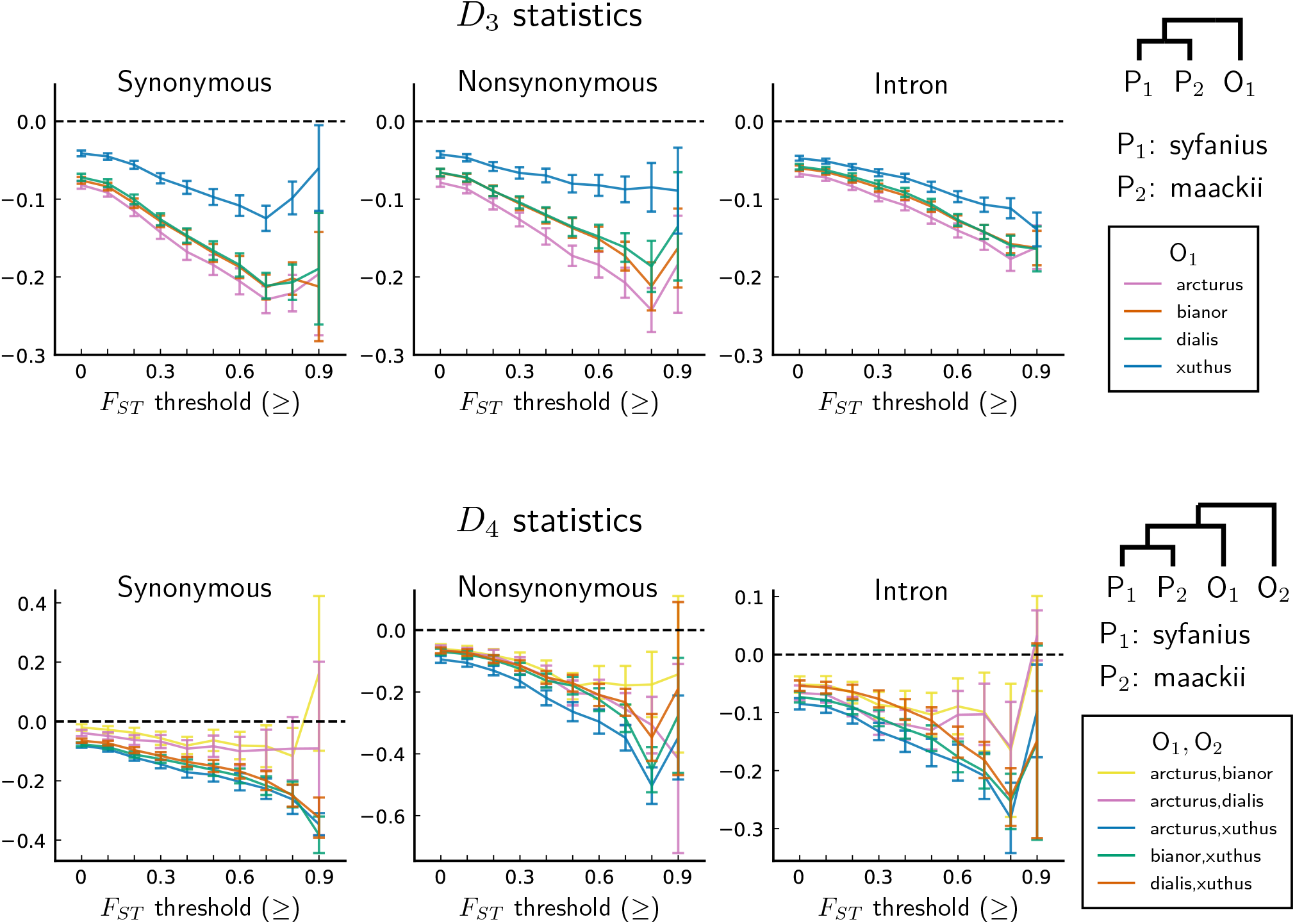
*D* statistics are unanimously negative. For each data point, we choose an *F*_*ST*_ threshold (x-axis) and report *D* statistics on SNPs with a background *F*_*ST*_ (50kb windows & 10kb increments) no less than the given threshold. Error bars are standard errors estimated using block-jackknife with 1Mb blocks.

**Figure 5:**
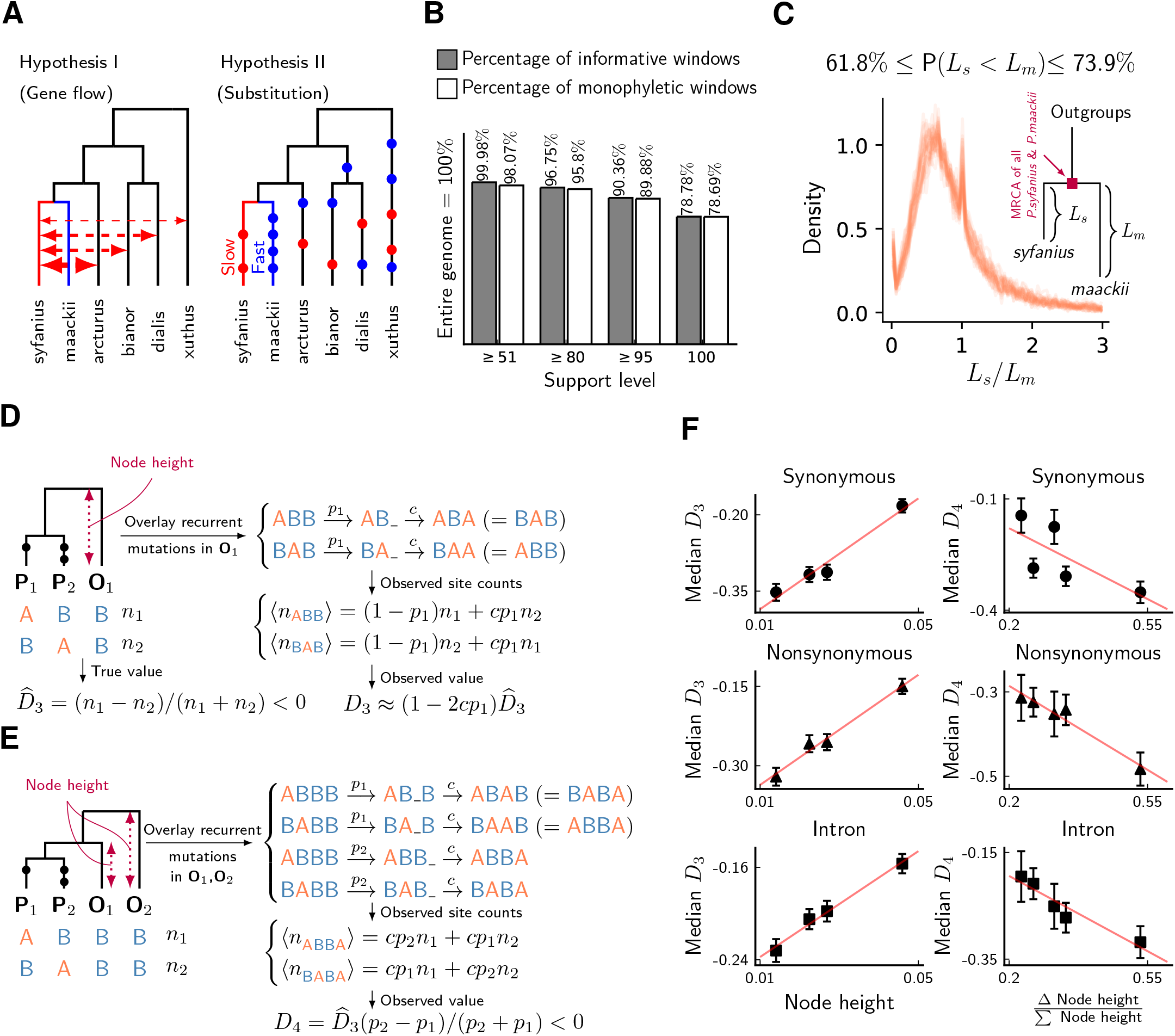
Unequal substitution rates between pure population of *P. syfanius* and *P. maackii*. **(A)** Two hypotheses to explain negative *D* statistics. **(B)** The percentage of local gene trees (50kb non-overlapping windows) where *P. syfanius* and *P. maackii* are together monophyletic. For each level of support, we filter out windows with *Support*(monophyly) and *Support*(paraphyly) below a given level, and report both the percentage of windows passing the filter (informative windows) and the percentage of monophyletic windows. **(C)** The distribution of *P. syfanius* branch lengths (*L_s_*) relative to those of *P. maackii* (*L_m_*) among highly supported monophyletic trees (*Support* 95). Each curve corresponds to a pairwise comparison between a *syfanius* individual and a *maackii* individual. Branch lengths are distances from tips to the most-recent common ancestor of all *syfanius*+*maackii* individuals. **(D)** Behavior of *D*_3_ under divergent substitution rates and recurrent mutations in outgroups (O_1_). **(E)** Behavior of *D*_4_ under divergent substitution rates and recurrent mutations in outgroups (O_1_ & O_2_). **(F)** Left: Median *D*_3_ is positively correlated with node height; Right: Median *D*_4_ is negatively correlated with ∆nodeheight/Σnodeheight. Node height is used as a proxy for the probability of recurrent mutation (*p_i_*)

### 2.4 Hybridization with outgroups does not explain site-pattern asymmetry

We first test hypothesis I using phylogenetic reconstruction using SNPs in annotated regions. We construct local gene trees for each 50kb non-overlapping window for all samples, including the four outgroup species. As biased gene flow with outgroups should rupture the monophyletic relationship among all *P. syfanius*+*P. maackii* individuals, the fraction of windows producing paraphyletic gene trees can be used to assess the potential impact of gene flow. However, almost all gene trees show the expected monophyletic relationship (Fig. 5B). This conclusion is independent of the level of support used to filter out genomic windows with ambiguous topologies (see Materials and Methods). Consequently, most windows show no phylogenetic signal of hybridization with outgroups. Nonetheless, among highly supported monophyletic trees (bootstrap support ≥ 95), *P. maackii* (in populations YA, BJ, XY) is always significantly more distant than *P. syfanius* from the most recent common ancestor of *P. maackii*+*P. syfanius* (Fig. 5C, significance levels reported via a Wilcoxon ranksum test for each pair of individuals, see Table S3). Second, the direction of allele sharing in the *D* statistics is unanimously biased towards *P. syfanius*. If hypothesis I were true, hybridization with outgroups is required to take place mainly in the highland lineage. There is no evidence to support why the highland lineage should receive more gene flow, as outgroups *P. xuthus* and *P. bianor* overlap broadly with both *P. maackii* and *P. syfanius*, while outgroup *P. dialis* is sympatric only with *P. maackii* (Condamine et al., 2013). Overall, we find little evidence for biased hybridization required by hypothesis I.

One might worry that by rejecting hypothesis I, we also throw doubt on widely accepted conclusions of the ABBA-BABA test for gene flow in other systems (that a significantly nonzero *D*_4_ implies hybridization with outgroups) (Durand et al., 2011). However, in the next section we show why *D*_3_ and *D*_4_ are fully consistent with hypothesis II, and so this is just a special case where the ABBA-BABA test produces a false positive for gene flow.

### 2.5 Evidence for divergent substitution rates

In hypothesis II, divergent substitution rates between *P. maackii* and *P. syfanius* interact with recurrent mutations in outgroups to produce asymmetric site patterns. To understand its effect on *D* statistics, consider a simplified model of recurrent mutation (Fig. 5D & 5E), where a site in outgroup *i* mutates with probability *p_i_*, producing the same derived allele with probability *c*. When averaged across the genome, *c* can be treated as a constant, and *p_i_* increases with distance to the outgroup. In the absence of gene flow, for three-taxon patterns, recurrent mutations modify *D*_3_ by a factor of approximately (1 − 2*cp*_1_) (Fig. 5D, see Materials and Methods), and observed *D*_3_ will thus be positively correlated with *p*_1_. For four-taxon patterns, it can be shown that observed *D*_4_ is always negative due to larger probabilities of recurrent mutation in more distant outgroups (Fig. 5E, see Materials and Methods). Assuming no significant contribution of incomplete lineage sorting (Maddison and Knowles, 2006), the value of *D*_4_ becomes more negative with increasing ∆*p_i_*/Σ*p_i_* = (*p*_2_ − *p*_1_)/(*p*_2_ + *p*_1_).

To test these signatures, we employ estimated node heights of outgroups in the mitochondrial tree (Fig. 2C) as proxies for outgroup distance, and hence for the relative probability of recurrent mutation (*p_i_*). In line with expected signatures, we find that observed *D*_3_ indeed increases with node height (Fig. 5F, left), and observed *D*_4_ decreases with (∆ Node height/∑ Node height) (Fig. 5F, right). Thus, the directions and magnitudes of both *D* statistics are congruent with hypothesis II. As hypothesis II naturally predicts unanimously negative *D*_3_ and *D*_4_ as well as their relative magnitudes among different outgroup combinations, it is more parsimonious than hypothesis I. Hence, divergent substitution rates likely exist between *P. syfanius* and *P. maackii*.

### 2.6 Rate-mixing at migration-drift equilibrium

Having established the existence of divergent substitution rates, we now explore how they become coupled by gene flow using a coalescent framework. As gene flow is ongoing between the two lineages, consider two haploid populations of size *N* exchanging genes at rate *m* (Fig. 6A). This simple isolation-with-migration (IM) model at equilibrium has relative divergence (Notohara, 1990)

**Figure 6:**
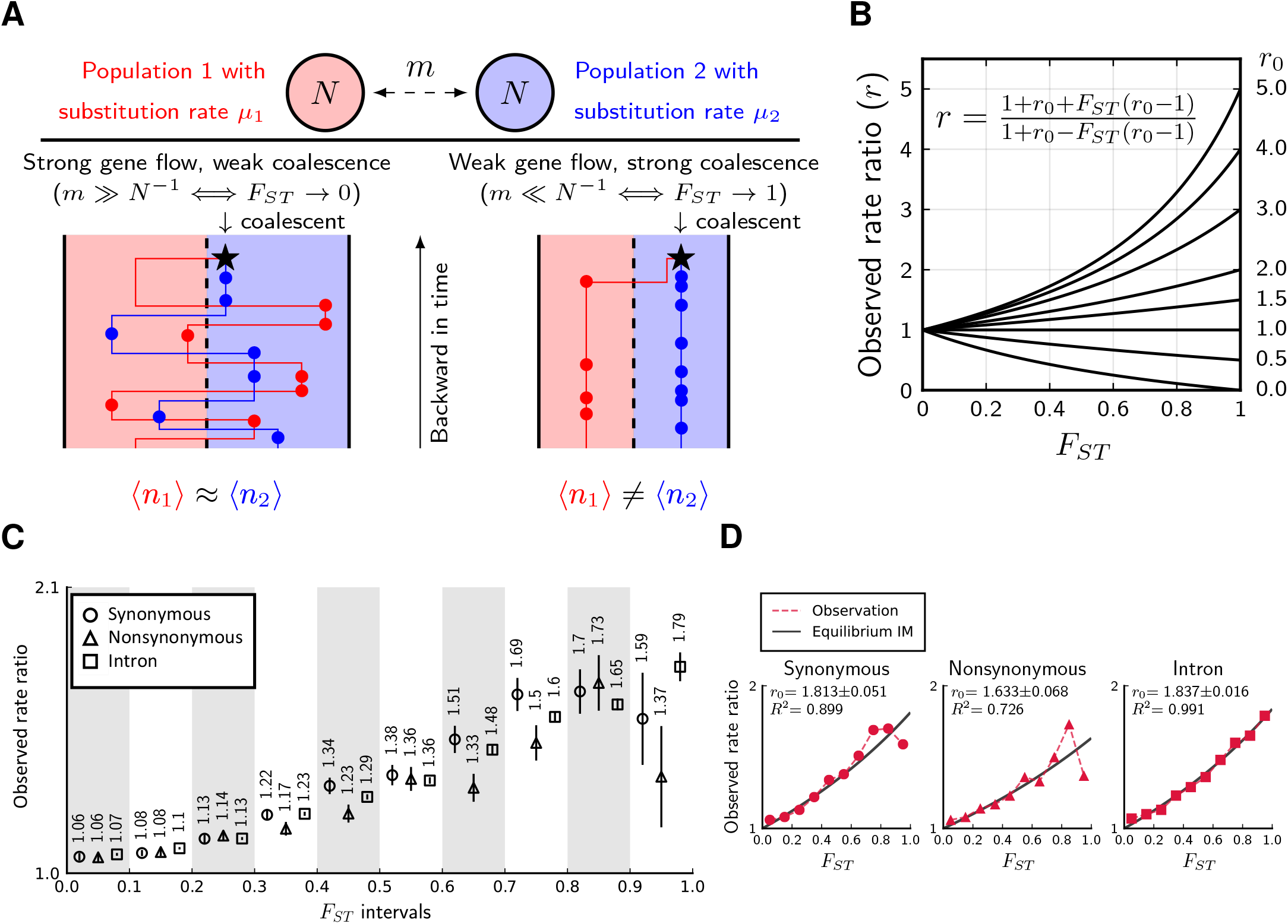
Divergence is correlated with increased differences in the relative number of substitutions. **(A)** Behavior of the equilibrium isolation-with-migration model with divergent substitution rates. If coalescence is weaker than gene flow, each lineage has a similar number of derived alleles. If coalescence is stronger than gene flow, lineages sampled from the population with a faster substitution rate will also inherit more derived alleles. **(B)** Theoretical relationship between observed rate ratio (*r*) and relative divergence (*F*_*ST*_), parameterized by the true ratio (*r*_0_) of substitution rates. **(C)** Observed rate ratios between *P. maackii* (population XY) and *P. syfanius* (Population KM), partitioned by ten *F*_*ST*_ intervals. Error bars are standard errors calculated using 1Mb block-jackknifing. **(D)** The theoretical relationship between *r* and *F*_*ST*_ is a good fit to observation.

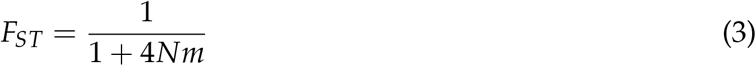

To quantify the signature of coupled rates, consider the asymmetry in observed numbers of substitutions (circles in Fig. 6A). Take a pair of sequences from two populations. Let ⟨*n*_1_⟩ be the expected number of derived alleles exclusive to the sequence in population 1, and let ⟨*n*_2_⟩ be the same expected number in population 2. Their ratio is defined as observed rate ratio:

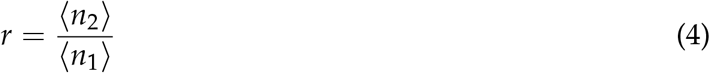

Evidently, *r* = 1 is the symmetric point where both sequences have the same number of derived alleles. Further, let the actual substitution rate in population 1 be *μ*_1_, and the actual substitution rate in population 2 be *μ*_2_. The ratio between the two actual rates are defined as the true rate ratio:

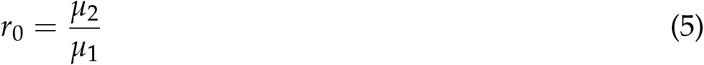

At migration-drift equilibrium, observed rate ratio (*r*) and observed divergence (*F*_*ST*_) are related by the following formula parameterized singly by *r*_0_ (Fig. 6B, see Materials and Methods):

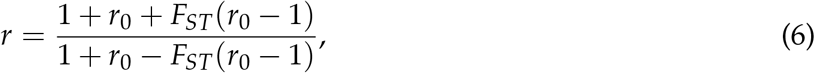

which translates into

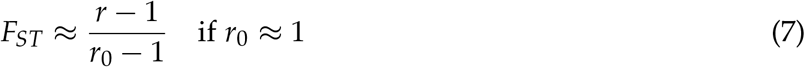

These formulae indicate that unequal substitution rates are more mixed in regions with lower genomic divergence. Eq. 7 is surprising, because it reveals that the remaining fraction of substitution rate divergence ((*r* − 1)/(*r*_0_ − 1)) is almost the same as *F*_*ST*_, which corresponds to the fraction of genetic variance explained by population structure (Wright, 1949). In Fig. 6B, the relationship between *r* and *F*_*ST*_ is still largely linear when one species evolves three times as fast as its sister species (*r*_0_ = 3), and so Eq. 7 might be robust under biologically realistic rates of substitutions between incipient species. Using extensive simulations (Fig. S4), we show that the full formula (Eq. 6) is also robust in a number of equilibrium population structures.

To test whether such predictions are met, we calculate *r* on synonymous, nonsynonymous, and intronic sites partitioned by their local *F*_*ST*_ values between pure populations (KM & XY), and recover a similar monotonic relationship across most *F*_*ST*_ partitions (Fig. 6C). For introns, the observed relationship between *r* and *F*_*ST*_ is a near perfect fit to Eq. 6 (Fig. 6D, squares), with an estimated *r*_0_ = 1.837. For synonymous (Fig. 6D, circles) and nonsynonymous sites (Fig. 6D, triangles), Eq. 6 also provides an excellent fit in regions with low to intermediate *F*_*ST*_. Estimated *r*_0_ for synonymous sites is 1.813, close to that of introns, while nonsynonymous sites have a considerably lower *r*_0_ of 1.633. If introns and synonymous substitutions are approximately neutral, we infer that neutral substitution rates are about 80% greater in the lowland species.

## 3. Discussion

In the gray zone of incomplete speciation, interspecific hybrids bridge between gene pools of divergent lineages (Mallet, 2005). We here demonstrate a similar role of hybridization in coupling and mixing differing substitution rates. Divergent rates of substitution carry information about outgroups, while divergence based on allele frequency differences does not (Barton, 1979). We emphasize that preservation of divergent substitution rates is a stronger effect than maintenance of allele frequency differences, because divergence of allele frequencies is a prerequisite for rate preservation. The dependence of the latter on the former can be coarsely quantified across the genome by the relationship between observed rate ratio *r* and relative divergence *F*_*ST*_ in an equilibrium system of hybridizing populations (Eq. 6). At migration-drift equilibrium, it is not surprising that divergent substitution rates are associated with relative divergence. In Fig. 6A, when coalescence occurs rapidly compared to gene flow, most substitutions separating individuals are species-specific. However, when gene flow is faster than coalescence, individuals will carry substitutions that occurred in both species. This could have important implications, because preserving lineage-specific substitution rates as measured by *r* might not require low absolute rates of gene flow. Instead, reducing effective population sizes via recurrent linked selection might achieve a similar result in populations at equilibrium (*N* ↓⇒ *F*_*ST*_ ↑).

Across non-equilibrium hybrid zones formed by secondary contact, reducing the absolute rate of gene flow by barrier loci in principle also keeps divergent rates from mixing, simply because it prevents substitutions accumulated in the allopatric phase from flowing between species. Interestingly, a higher substitution rate in the lowland lineage *P. maackii* is congruent with the evolutionary speed hypothesis (Rensch, 1959), where evolution accelerates in warmer climates. It is unclear what mechanisms generate increased substitution rates, but the lowland lineage typically has an additional autumn brood that is absent in *P. syfanius* (Takasaki et al., 2007) (Fig. S1F). Warmer temperatures in lowland habitats might also increase spontaneous mutation rates in ectothermic insects (Waldvogel and Pfenninger, 2021).

An additional result from our study is that divergent substitution rates might produce spuriously non-zero *D*_4_ statistics when combined with recurrent mutations, which could increase the false positive rate of the ABBA-BABA test. This phenomenon has been suspected in humans (Amos, 2020), and is certainly a theoretical possibility (Hibbins and Hahn, 2022), but has not been tested in most empirical studies.

## 4. Materials and Methods

### 4.1 Museum specimens and climate Data

Museum specimens with verifiable locality data of all species were gathered from The University Museum (The University of Tokyo), Global Biodiversity Information Facility, and individual collectors (Table S1). Records of *P. maackii* from Japan, Korea and NE China were excluded from the analysis, so that most *P. maackii* individuals correspond to *ssp. shimogorii*, the subspecies that hybridizes with *P. syfanius*. Spatial principal component analysis was performed on elevation, maximum temperature of warmest month, minimum temperature of coldest month, and annual precipitation, all with 30s resolution from WorldClim-2 (Fick and Hijmans, 2017). The first two PCAs, combined with tree cover (Hansen et al., 2013), were used in MaxEnt-3.4.1 to produce species distribution models that use known localities to predict occurrence probabilities across the entire landscape (Phillips et al., 2017). Outputs were trimmed near known boundaries of each species. See Fig. S1B for the final result.

### 4.2 Sampling, re-sequencing, and mitochondrial phylogeny

Eleven males of *P. syfanius* and *P. maackii*, with one male of *P. arcturus* and one male of *P. dialis* were collected in the field between July and August in 2018 (Table S1), and were stored in RNAlater at −20C prior to DNA extraction. E.Z.N.A Tissue DNA kit was used to extract genomic DNA, and KAPA DNA HyperPlus 1/4 was used for library preparation, with an insert size of 350bp and 2 PCR cycles. The library is sequenced on a Illumina NovaSeq machine with paired-end reads of 150bp. Adaptors were trimmed using Cutadapt-1.8.1, and subsequently the reads were mapped to the reference genome of *P. bianor* with BWA-0.7.15, then deduplicated and sorted via PicardTools-2.9.0. The average coverage among 13 individuals in non-repetitive regions varies between 20× to 30×. Variants were called twice using BCFtools-1.9 – the first including all samples, used in analyses involving outgroups, and the second excluding *P. arcturus* and *P. dialis*, used in all other analyses.

The following thresholds were used to filter variants: 10*N* <DP < 50*N*, where *N* is the sample size; QUAL> 30; MQ> 40; MQ0F < 0.2. As a comparison, we also called variants with GATK4 and followed its best practices, and 93% of post-filtered SNPs called by GATK4 overlapped with those called by BCFtools. We used SNPs called by BCFtools throughout the analysis. Mitochondrial genomes were assembled from trimmed reads with NOVOPlasty-4.3.1 (Dierckxsens et al., 2017), using a published mitochondrial ND5 gene sequence of *P. maackii* as a bait (NCBI accession number: AB239823.1). We also used the following published mitochondrial genomes (NCBI accession numbers): KR822739.1, NC_029244.1, JN019809.1. The neighbor-joining mitochondrial phylogeny was built with Geneious Prime-2021.2.2 (genetic distance model: Tamura-Nei), and we used 104 replicates for bootstrapping. The reference genome of *P. xuthus* was previously aligned to the genome of *P. bianor* and we used this alignment directly in all analysis (Lu et al., 2019).

### 4.3 Calculating site-pattern asymmetry

Given a species tree {{P_1_,P_2_},O}, where P_1_ and P_2_ are sister species and O is an outgroup, if mutation rates are equal between {P_1_,P_2_}, and no gene flow with O, then on average the number of derived alleles in P_1_ should equal the number of derived alleles in P_2_. Let *S* be a collection of sites, *f_s_* be the frequency of a particular site pattern at site *s* ∈ *S*. “ABB” be the pattern where only P_2_ and O share the same allele, and “BAB” be the pattern where only P_1_ and O share the same allele, then the three-species *D*_3_ statistic is calculated as

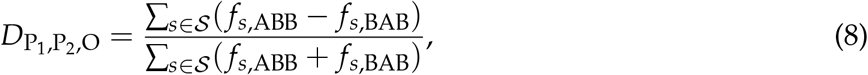

where *S* is always limited to sites without polymorphism in the outgroup O. This statistic is in principle capturing the same source of asymmetry as the statistic proposed by Hahn & Hibbins (Hahn and Hibbins, 2019), although their version uses divergence to the outgroup instead of frequencies of site-counts. Similarly, the four-species *D*_4_ statistic, which considers species tree {{{P_1_,P_2_},O_1_},O_2_} and site patterns ABBA versus BABA (Durand et al., 2011) is calculated as

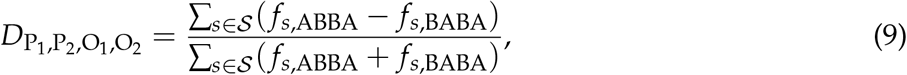

where *S* is always limited to sites without polymorphism in the second outgroup O_2_. The significance of both tests was computed using block-jackknife over 1Mb blocks across the genome. Additionally, we estimated rate ratio as follows. First we restricted to sites where all outgroups are fixed for the same ancestral allele to dampen the influence of recurrent mutation. Then, for each site, sample one allele at random from each focal lineage. Calculate the probability of observing a derived allele in P_1_ but not in P_2_, and the probability of observing a derived allele in P_2_ but not in P_1_. The rate ratio is computed as the ratio between the two probabilities. Explicitly, let *I*(·) be the identity function, and *f_s_* be the frequency of the derived allele, then:

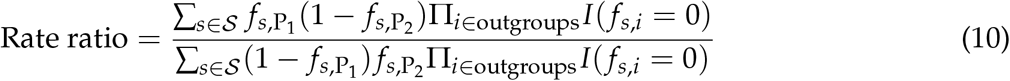

Its standard error was estimated using 1Mb block-jackknifing. We excluded *P. xuthus* from the outgroups to increase the number of informative sites when using this formula.

### 4.4 *D*_3_ and *D*_4_ under unequal substitution rates and recurrent mutations

In this section, we calculate observed *D*_3_ and *D*_4_ assuming that incomplete lineage sorting contributes insignificantly to both statistics. If incomplete lineage sorting is present, it will not create new bias (numerators are on average unchanged), but will likely dampen existing bias (inflating denominators).

As substitutions are independent along each lineage, we can mute recurrent mutations in outgroups and generate them afterwards. For three taxa with gene tree {{P_1_,P_2_},O_1_}, before recurrent mutations, there are *n*_1_ sites with pattern (B,A,A), and *n*_2_ sites with pattern (A,B,A). If substitution rate is higher in P_2_, we have *n*_2_ > *n*_1_, so the true value of *D*_3_, written as 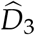, is always negative:

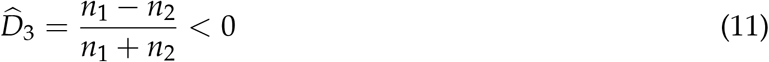

Next, recurrent mutations in O_1_ occur at each site with an average probability *p*_1_, and with an average probability *c*, ancestral alleles from affected sites in O_1_ are converted to the same derived allele in {P_1_,P_2_}. *c* will be independent of *n*_1_ and *n*_2_, as long as substitutions between {P_1_,P_2_} are only different in *rates*, but not mutation types. Hence, two possible mutation paths exist:

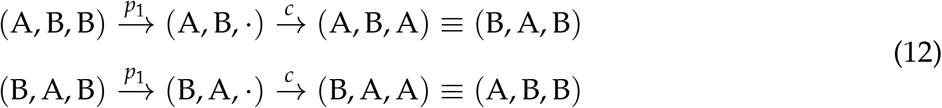

The expected site counts, after recurrent mutations, become

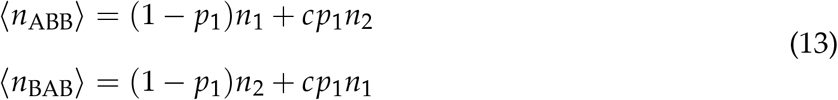

Using the new expected site counts in *D*_3_ statistics produce the following value:

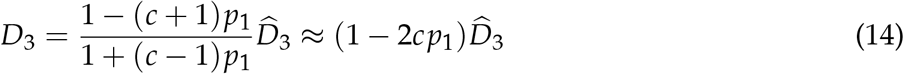

Since 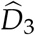 is negative, it grows approximately linearly near small values of *p*_1_. (The full equation is still monotonic in *p*_1_.)

Similarly, for four-taxon statistics, before recurrent mutation, there are two types of sites: (A,B,B,B)-*n*_1_; (B,A,B,B)-*n*_2_. Suppose the average probability of recurrent mutation is *p*_1_ in O_1_, and *p*_2_ in O_2_, and the conversion probability of each recurrent mutation into derived alleles of {P_1_, P_2_} is *c* for both outgroups. Using the same procedure, one can show that

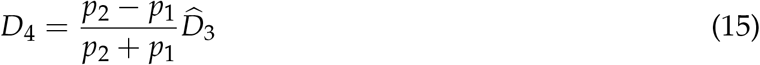

Since recurrent mutations occur more frequently in distant outgroups, *p*_2_ > *p*_1_. Because 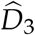 is negative, we have *D*_4_ < 0.

### 4.5 Local gene trees

Local gene trees were estimated using iqtree-2.0 (Minh et al., 2020) on 50kb non-overlapping genomic windows with options -`m MFP -B 5000`. Only SNPs from annotated regions (synonymous sites + nonsynonymous sites + introns) across all individuals were used. For diploid individuals, heterozygous sites were assigned IUPAC ambiguity codes and iqtree assigned equal likelihood for each underlying character, thus information from heterozygous sites is largely retained. This is crucial as we are interested in the branch length of inferred trees. Option -`m MFP` implements iqtree’s ModelFinder that tests the FreeRate model to accommodate maximum flexibility with rate-variation among sites (Kalyaanamoorthy et al., 2017). We also used UltraFast Bootstrap to calculate the support for different types of splits in each window (the -`B` 5000 option) (Hoang et al., 2018). In each window, we extracted the support for monophyly among *P. maackii*+*P.syfanius* directly from the output of UltraFast Bootstrap, and we define the support for paraphyly among *P. maackii*+*P.syfanius* as (100 - the support for monophyly). For each level of support, we filtered out genomic windows where both the support for monophyly and the support for paraphyly drop below the given level. The remaining windows were considered informative.

### 4.6 Rate-mixing under the equilibrium IM model

We construct a continuous-time coalescent model as follows. Both populations have *N* haploid individuals, gene flow rate is *m*, and coalescent rate is *N*−1 in each population. In the equilibrium system, as we track both haploid individuals backward in time, there are six distinct states: (1|2), (2|1), (1, 2|), (|1, 2), (0|), (|0), where 1 & 2 represent two individuals prior to coalescent, 0 is the state of coalescent, and (·|·) shows the location of each lineage. Its transition density p(*t*) satisfies *∂_t_***p** = **Ap**, where **A** is given as

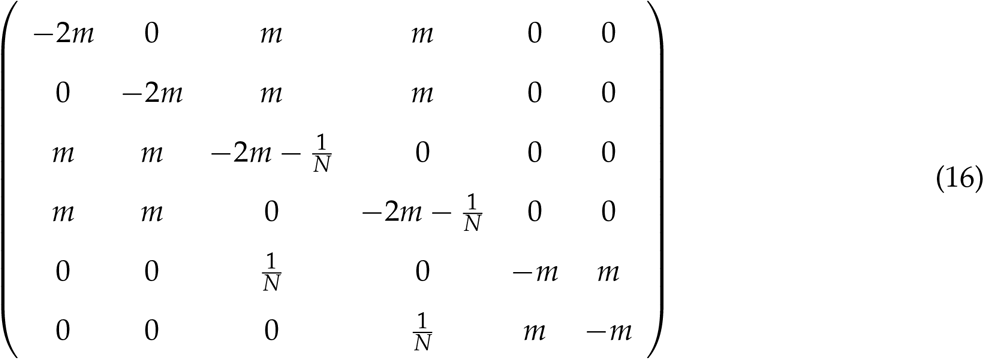

Let *S_i_*_|*j*_(*T*) be the mean sojourn time of an uncoalesced individual inside population *i* during 0 ≤ *t* ≤ *T*, conditioning on the individual being taken from population *j* at *t* = 0. Assuming the infinite-site mutation model, let *μ_i_* be the substitution rate in population *i*, observed rate ratio *r* is thus

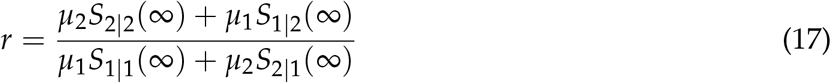

where (due to symmetry)

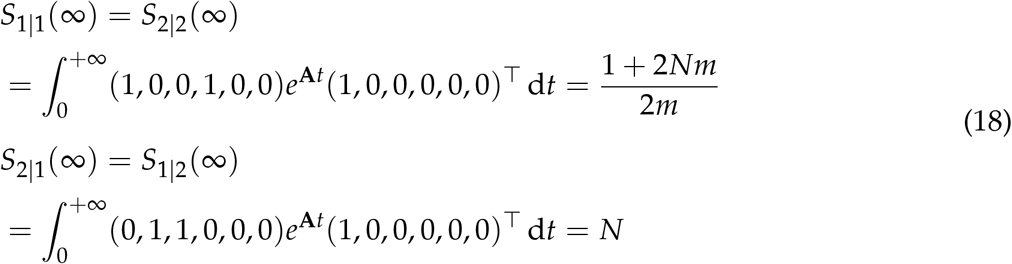

Let *r*_0_ = *μ*_2_/*μ*_1_, and since *F*_*ST*_ = (1 + 4*Nm*)^−1^, we have

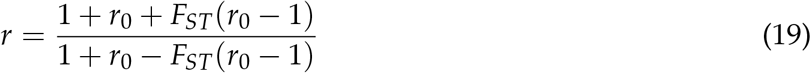

### 4.7 Local ancestry estimation

Software ELAI (Guan, 2014) with a double-layer HMM model was used to estimate diploid local ancestries across chromosomes. An example command is as follows:

`elai-lin -g genotype.maackii.txt -p 10 -g genotype.syfanius.txt -p 11 -g genotype.admixed.txt -p 1 -pos position.txt -s 30 -C 2 -c 10 -mg 5000 -exclude-nopos`. Note that -`mg` specifies the resolution of ancestry blocks, thus increasing its value will increase the stochastic error of incorrectly inferring very short blocks of ancestry. To control for uncertainty, we estimated repeatedly for 50 times. All replicates were used simultaneously in finding the correlation coefficients between entropy and other variables. Results from an example run is in Fig. S5.

### 4.8 Ancestry and entropy

Here we introduce concisely the data transformation framework for calculating the entropy of local ancestry. The mathematical detail of this approach is presented in Supplementary Information Section 1.

#### Ancestry representation

The space of all ancestry signals is high-dimensional, and directly calculating the entropy in this space is not feasible with just a few individuals. So we propose to measure only the pairwise correlation of ancestries among sites, which captures only the second-order randomness, but is sufficient for practical purposes. Consider a hybrid individual with two parental populations indexed by *k* = 1, 2. Assuming a continuous genome, let *p_k_*(*l*) = 0, 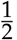, 1 be the diploid ancestry of locus *l* within genomic interval [0, *L*]. By definition, we have *p*_1_(*l*) + *p*_2_(*l*) = 1, i.e. the total ancestry is conserved everywhere in the genome. The bi-ancestry signal at locus *l* is defined as the following complex variable

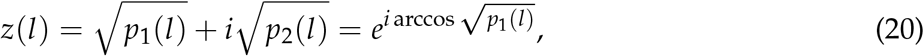

where 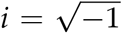 is the imaginary number. An advantage of using a complex representation for the bi-ancestry signal is that we can model different ancestries along the genome as different phases of a complex unit phasor (*e*^*iθ*^), such that the power of the signal at any given locus is simply the sum of both ancestries, which is conserved (|*z*(*l*)|^2^ = 1). It ensures that we do not bias the analysis to any particular region or any particular individual when decomposing the signal into its spectral components.

#### Within-individual spectral entropy (S_w_)

To characterize the average autocorrelation along a an ancestry signal at a given scale *l*, define the following scale-dependent autocorrelation function 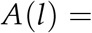 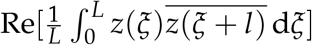, where *z*(*l*) is understood as a periodic function such that *z*(*ξ* + *l*) = *z*(*ξ* + *l* − *L*) whenever the position goes outside of [0, *L*]. The Wiener-Khinchin theorem guarantees that *z*(*l*)’s power spectrum *ζ*(*f*), which is discrete, and the autocorrelation function *A*(*l*) form a Fourier-transform pair. Due to the uncertainty principle of Fourier transform, *A*(*l*) that vanishes quickly at short distances (small-scale autocorrelation) will produce a wide *ζ*(*f*), and vice versa. So the entropy *S*_*w*_ of *ζ*(*f*), which measures the spread of the total ancestry into each spectral component, also measures the scale of autocorrelation. In practice, *ζ*(*f*) is the square modulus of the Fourier series coefficients of *z*(*l*), and we fold the spectrum around *f* = 0 before calculating the within-individual entropy *S*_*w*_. The formula used in the manuscript is

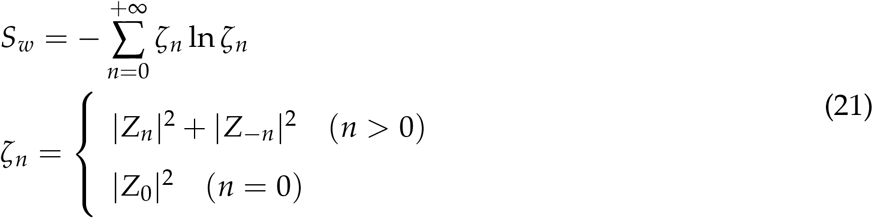

where *Z_n_* are the Fourier coefficients from the expansion 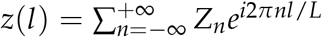. To speed up the Fourier expansion, we could *densely* pack equally-spaced markers that sample a continuous ancestry signal into a discrete signal, which then undergoes Fast Fourier Transform (FFT). The spectrum of FFT (discrete and finite) approximates the continuous-time Fourier spectrum (discrete and infinite), and entropy also converges as marker density increases.

#### Between-individual spectral entropy (S_b_)

As ancestry configuration is far from random around barrier loci, it will also influence the correlation of ancestry between different individuals at the same locus. For a genomic region experiencing strong barrier effects, two individuals could either be very similar in ancestry, or very different. This effect can be quantified by first calculating the crosscorrelation 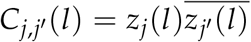 at position *l* between individuals *j* and *j*′, and then averaging across a genome interval: 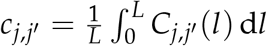. The *J* × *J* dimensional matrix **C** with entries *c*_*j*,*j*_′ describes the pairwise cross-correlation within the cohort of *J* individuals. We also have *c_j_*_,*j*_ ≡ 1 as each individual is perfectly correlated with itself. The matrix **C** is Hermitian, so it has a real spectral decomposition with eigenvalues *λ_j_* that satisfy ∑_*j*_ λ_*j*_/*J* = 1. This process is very similar to performing a principal component analysis on the entire cohort of individuals, and *λ_j_*/*J* describes the fraction of the total ancestry projected onto principal component *j*. If many loci co-vary in ancestry, the spectrum {*λ_j_*} will be concentrated near the first few components. Similarly, we use entropy to measure the spread of the spectrum, and hence the between-individual spectral entropy is defined as

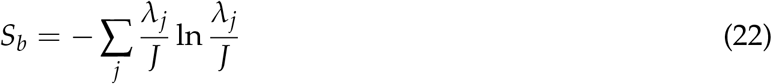

## 5. Data Availability

Relevant code is available at: https://github.com/tzxiong/2021_Maackii_Syfanius_HybridZone

Whole-genome sequences are deposited in the National Center for Biotechnology Information, Sequence Read Archive (BioProject Accession Number: PRJNA765117).

## 6. Author Contributions

T.X. and J.M. designed the project. X.L. provided reference genomes for *P. bianor* and *P. xuthus*, and also facilitated the fieldwork. M.Y. provided museum specimens used in the manuscript. T.X. collected and analyzed the samples. T.X. and J.M. wrote the manuscript.

## 7. Acknowledgement

This work was supported by the Lewis and Clark Fund for Exploration and Field Research (American Philosophical Society, 2017-2018) granted to T.X.; T.X. is also funded by a studentship from Harvard Department of Organismic and Evolutionary Biology, the NSF-Simons Center for Mathematical and Statistical Analysis of Biology at Harvard (award number #1764269) and the Harvard Quantitative Biology Initiative during the project. We thank Harvard FAS Research Computing for providing computation resources. We thank Fernando Seixas for discussion on hybrid ancestry inference. We thank Kaifeng Bu for his comments on information-theoretic measures. We thank Nathaniel Edelman, Neil Rosser, Miriam Miyagi, Yuttapong Thawornwattana, Sarah Dendy, Liang Qiao, John Wakeley, Robin Hopkins, and Naomi Pierce for their valuable input. We also thank Yuchen Zheng, Zhuoheng Jiang, Shaoji Hu, Feng Cao, and Shui Xu for support in the field. Feng Cao provided specimen photos from the hybrid zone.

## References

Amos, W. (2020). Signals interpreted as archaic introgression appear to be driven primarily by faster evolution in Africa. Royal Society Open Science, 7(7):191900.

Arbeithuber, B., Betancourt, A. J., Ebner, T., and Tiemann-Boege, I. (2015). Crossovers are associated with mutation and biased gene conversion at recombination hotspots. Proceedings of the National Academy of Sciences, 112(7):2109–2114.

Barton, N. H. (1979). Gene flow past a cline. Heredity, 43(3):333–339.

Barton, N. H. (1983). Multilocus clines. Evolution, pages 454–471.

Bromham, L. and Penny, D. (2003). The modern molecular clock. Nature Reviews Genetics, 4(3):216–224.

Condamine, F. L., Toussaint, E. F., Cotton, A. M., Genson, G. S., Sperling, F. A., and Kergoat, G. J. (2013). Fine-scale biogeographical and temporal diversification processes of peacock swallowtails (*Papilio* subgenus *Achillides*) in the Indo-Australian Archipelago. Cladistics, 29(1):88–111.

Costa, R. J. and Wilkinson-Herbots, H. (2017). Inference of gene flow in the process of speciation: An efficient maximum-likelihood method for the isolation-with-initial-migration model. Genetics, 205(4):1597–1618.

DeWitt, W. S., Harris, K. D., Ragsdale, A. P., and Harris, K. (2021). Nonparametric coalescent inference of mutation spectrum history and demography. Proceedings of the National Academy of Sciences, 118(21).

Dierckxsens, N., Mardulyn, P., and Smits, G. (2017). Novoplasty: de novo assembly of organelle genomes from whole genome data. Nucleic Acids Research, 45(4):e18–e18.

Durand, E. Y., Patterson, N., Reich, D., and Slatkin, M. (2011). Testing for ancient admixture between closely related populations. Molecular Biology and Evolution, 28(8):2239–2252.

Fick, S. E. and Hijmans, R. J. (2017). WorldClim 2: new 1-km spatial resolution climate surfaces for global land areas. International Journal of Climatology, 37(12):4302–4315.

Guan, Y. (2014). Detecting structure of haplotypes and local ancestry. Genetics, 196(3):625–642.

Hahn, M. W. and Hibbins, M. S. (2019). A three-sample test for introgression. Molecular Biology and Evolution, 36(12):2878–2882.

Hansen, M. C., Potapov, P. V., Moore, R., Hancher, M., Turubanova, S. A., Tyukavina, A., Thau, D., Stehman, S. V., Goetz, S. J., Loveland, T. R., et al. (2013). High-resolution global maps of 21st century forest cover change. Science, 342(6160):850–853.

Harris, K. (2015). Evidence for recent, population-specific evolution of the human mutation rate. Proceedings of the National Academy of Sciences, 112(11):3439–3444.

Hasselman, D. J., Argo, E. E., McBride, M. C., Bentzen, P., Schultz, T. F., Perez-Umphrey, A. A., and Palkovacs, E. P. (2014). Human disturbance causes the formation of a hybrid swarm between two naturally sympatric fish species. Molecular Ecology, 23(5):1137–1152.

Hibbins, M. S. and Hahn, M. W. (2022). Phylogenomic approaches to detecting and characterizing introgression. Genetics, 220(2):iyab173.

Hoang, D. T., Chernomor, O., Von Haeseler, A., Minh, B. Q., and Vinh, L. S. (2018). UFBoot2: improving the ultrafast bootstrap approximation. Molecular Biology and Evolution, 35(2):518–522.

Hudson, R. R. (1990). Gene genealogies and the coalescent process. Oxford Surveys in Evolutionary Biology, 7(1):44.

Irwin, D. E., Milá, B., Toews, D. P., Brelsford, A., Kenyon, H. L., Porter, A. N., Grossen, C., Delmore, K. E., Alcaide, M., and Irwin, J. H. (2018). A comparison of genomic islands of differentiation across three young avian species pairs. Molecular Ecology, 27(23):4839–4855.

Jensen-Seaman, M. I., Furey, T. S., Payseur, B. A., Lu, Y., Roskin, K. M., Chen, C.-F, Thomas, M. A., Haussler, D., and Jacob, H. J. (2004). Comparative recombination rates in the rat, mouse, and human genomes. Genome Research, 14(4):528–538.

Kalyaanamoorthy, S., Minh, B. Q., Wong, T. K., Von Haeseler, A., and Jermiin, L. S. (2017). ModelFinder: fast model selection for accurate phylogenetic estimates. Nature Methods, 14(6):587–589.

Kashiwabara, S. (1991). Why are *Papilio dehaanii* from Tokara Is. and Izu Is. beautiful? Choken-Field, 6(9):6–16.

Kautt, A. F., Kratochwil, C. F., Nater, A., Machado-Schiaffino, G., Olave, M., Henning, F., Torres-Dowdall, J., Härer, A., Hulsey, C. D., Franchini, P., et al. (2020). Contrasting signatures of genomic divergence during sympatric speciation. Nature, 588(7836):106–111.

Kronforst, M. R., Hansen, M. E., Crawford, N. G., Gallant, J. R., Zhang, W., Kulathinal, R. J., Ka-pan, D. D., and Mullen, S. P. (2013). Hybridization reveals the evolving genomic architecture of speciation. Cell Reports, 5(3):666–677.

Lepage, T., Bryant, D., Philippe, H., and Lartillot, N. (2007). A general comparison of relaxed molecular clock models. Molecular Biology and Evolution, 24(12):2669–2680.

Lercher, M. J. and Hurst, L. D. (2002). Human SNP variability and mutation rate are higher in regions of high recombination. Trends in Genetics, 18(7):337–340.

Li, X., Fan, D., Zhang, W., Liu, G., Zhang, L., Zhao, L., Fang, X., Chen, L., Dong, Y., Chen, Y., et al. (2015). Outbred genome sequencing and CRISPR/Cas9 gene editing in butterflies. Nature Communications, 6(1):1–10.

Liu, H., Jia, Y., Sun, X., Tian, D., Hurst, L. D., and Yang, S. (2016). Direct determination of the mutation rate in the bumblebee reveals evidence for weak recombination-associated mutation and an approximate rate constancy in insects. Molecular Biology and Evolution, 34(1):119–130.

Lu, S., Yang, J., Dai, X., Xie, F., He, J., Dong, Z., Mao, J., Liu, G., Chang, Z., Zhao, R., et al. (2019). Chromosomal-level reference genome of chinese peacock butterfly (*Papilio bianor*) based on thirdgeneration DNA sequencing and Hi-C analysis. GigaScience, 8(11):giz128.

Lynch, M. (2010). Evolution of the mutation rate. Trends in Genetics, 26(8):345–352.

Maddison, W. P. and Knowles, L. L. (2006). Inferring phylogeny despite incomplete lineage sorting. Systematic Biology, 55(1):21–30.

Mallet, J. (2005). Hybridization as an invasion of the genome. Trends in Ecology & Evolution, 20(5):229–237.

Michel, A. P., Sim, S., Powell, T. H., Taylor, M. S., Nosil, P., and Feder, J. L. (2010). Widespread genomic divergence during sympatric speciation. Proceedings of the National Academy of Sciences, 107(21):9724–9729.

Minh, B. Q., Schmidt, H. A., Chernomor, O., Schrempf, D., Woodhams, M. D., Von Haeseler, A., and Lanfear, R. (2020). IQ-TREE 2: new models and efficient methods for phylogenetic inference in the genomic era. Molecular Biology and Evolution, 37(5):1530–1534.

Nosil, P., Funk, D. J., and Ortiz-Barrientos, D. (2009). Divergent selection and heterogeneous genomic divergence. Molecular Ecology, 18(3):375–402.

Notohara, M. (1990). The coalescent and the genealogical process in geographically structured population. Journal of Mathematical Biology, 29(1):59–75.

Ohta, T. (1993). An examination of the generation-time effect on molecular evolution. Proceedings of the National Academy of Sciences, 90(22):10676–10680.

Payseur, B. A. and Rieseberg, L. H. (2016). A genomic perspective on hybridization and speciation. Molecular Ecology, 25(11):2337–2360.

Phillips, S. J., Anderson, R. P., Dudík, M., Schapire, R. E., and Blair, M. E. (2017). Opening the black box: An open-source release of Maxent. Ecography, 40(7):887–893.

Presgraves, D. C. (2018). Evaluating genomic signatures of “the large X-effect” during complex speciation. Molecular Ecology, 27(19):3822–3830.

Ravinet, M., Faria, R., Butlin, R., Galindo, J., Bierne, N., Rafajlović, M., Noor, M., Mehlig, B., and Westram, A. (2017). Interpreting the genomic landscape of speciation: a road map for finding barriers to gene flow. Journal of Evolutionary Biology, 30(8):1450–1477.

Renaut, S., Grassa, C., Yeaman, S., Moyers, B., Lai, Z., Kane, N., Bowers, J., Burke, J., and Rieseberg, L. (2013). Genomic islands of divergence are not affected by geography of speciation in sunflowers. Nature Communications, 4(1):1–8.

Rensch, B. (1959). Evolution above the species level. Columbia University Press.

Sedghifar, A., Brandvain, Y., and Ralph, P. (2016). Beyond clines: lineages and haplotype blocks in hybrid zones. Molecular Ecology, 25(11):2559–2576.

Takasaki, H., Kawaguchi, N., Kuribayashi, T., Hasuo, R., and Kobayashi, S. (2007). Unusual successive occurrence of the Maackii Peacock (*Papilio maackii*; Papilionidae) at Okayama University of Science, southwestern Honshu lowland, in 2006 summer and autumn. Naturalistae, (11):11–14.

Wakeley, J. (2016). Coalescent Theory: An Introduction. Macmillan Learning.

Waldvogel, A.-M and Pfenninger, M. (2021). Temperature dependence of spontaneous mutation rates. Genome Research, 31(9):1582–1589.

Wolf, J. B. and Ellegren, H. (2017). Making sense of genomic islands of differentiation in light of speciation. Nature Reviews Genetics, 18(2):87–100.

Wright, S. (1949). The genetical structure of populations. Annals of Eugenics, 15(1):323–354.

Yang, S., Wang, L., Huang, J., Zhang, X., Yuan, Y., Chen, J.-Q., Hurst, L. D., and Tian, D. (2015). Parent–progeny sequencing indicates higher mutation rates in heterozygotes. Nature, 523(7561):463–467.

Yang, Z. and Rannala, B. (2012). Molecular phylogenetics: principles and practice. Nature Reviews Genetics, 13(5):303–314.

