## Supplementary Information for "Admixture of evolutionary rates across a hybrid zone"

**Table S1:** Samples and localities

| **Non-genetic samples** |
|---|
| 1. The University Museum, The University of Tokyo (Harada et al., 2012; Yago et al., 2021). |
| 2. Global Biodiversity Information Facility (The Global Biodiversity Information Facility, 2021a,b). |

**Genetic samples**

| Sample name NCBI Accession | Current taxonomy name | Name in the manuscript | Locality | Collecting date | Sex |
|---|---|---|---|---|---|
| TXSC20180711006 SAMN21542929 | *Papilio maackii shimogorii* | *Papilio maackii* | XY | Jul-11-2018 | ♂ |
| TXSC20180711011 SAMN21542930 | *Papilio maackii shimogorii* | *Papilio maackii* | XY | Jul-11-2018 | ♂ |
| TXSC20180713001 SAMN21542931 | *Papilio maackii shimogorii* | *Papilio maackii*-hybrid | BJ | Jul-13-2018 | ♂ |
| TXSC20180727005 SAMN21542932 | *Papilio maackii shimogorii* | *Papilio maackii* | YA | Jul-27-2018 | ♂ |
| TXSC20180713007 SAMN21542933 | *Papilio maackii albosyfanius* | *Papilio syfanius*-hybrid | WN | Jul-13-2018 | ♂ |
| TXSC20180713008 SAMN21542934 | *Papilio maackii albosyfanius* | *Papilio syfanius*-hybrid | WN | Jul-13-2018 | ♂ |
| TXSC20180714006 SAMN21542935 | *Papilio maackii albosyfanius* | *Papilio syfanius*-hybrid | WN | Jul-14-2018 | ♂ |
| TXSC20180714008 SAMN21542936 | *Papilio maackii albosyfanius* | *Papilio syfanius*-hybrid | WN | Jul-14-2018 | ♂ |
| TXSC20180718001 SAMN21542937 | *Papilio maackii albosyfanius* | *Papilio syfanius* | KM | Jul-18-2018 | ♂ |
| TXSC20180718009 SAMN21542938 | *Papilio maackii albosyfanius* | *Papilio syfanius* | KM | Jul-18-2018 | ♂ |
| TXSC20180718016 SAMN21542939 | *Papilio maackii albosyfanius* | *Papilio syfanius* | KM | Jul-18-2018 | ♂ |
| TXSC20180714007 SAMN21542940 | *Papilio arcturus* | *Papilio arcturus* | WN | Jul-14-2018 | ♂ |
| TXSC20180821006 SAMN21542941 | *Papilio dialis* | *Papilio dialis* | NB | Aug-21-2018 | ♂ |
| Reference genome* | *Papilio bianor* | *Papilio bianor* | - | - | ♂ |
| Reference genome** | *Papilio xuthus* | *Papilio xuthus* | - | - | ♂ |

*The reference genome of *P. bianor* was assembled in (Lu et al., 2019), and was used for all analyses involving *P. bianor*. The reference genome of *P. bianor* used one adult male and two larval males from populations in Kunming City (KM).
** We used the alignment between *P. bianor* and *P. xuthus* in (Lu et al., 2019) for all analysis involving *P. xuthus*.

| Locality | Latitude | Longitude | Elevation | Detail |
|---|---|---|---|---|
| YA | 30.06169 | 102.98816 | 1157 | Bifeng Valley, Ya'an, Sichuan Province, China |
| XY | 28.29648 | 105.53281 | 329 | Huagaoxi, Xuyong County, Luzhou, Sichuan Province, China |
| BJ | 27.31812 | 105.27144 | 1743 | Wenbi Mountain, Bijie, Guizhou Province, China |
| WN | 26.75528 | 104.4317 | 2358 | Jinzhongzhen, Weining County, Bijie, Guizhou Province, China |
| KM | 25.08828 | 102.80779 | 2038 | Huanglongqing, Kunming, Yunnan Province, China |
| NB | 29.799100 | 121.252986 | 200 | Wulongtan, Ningbo, Zhejiang Province, China |

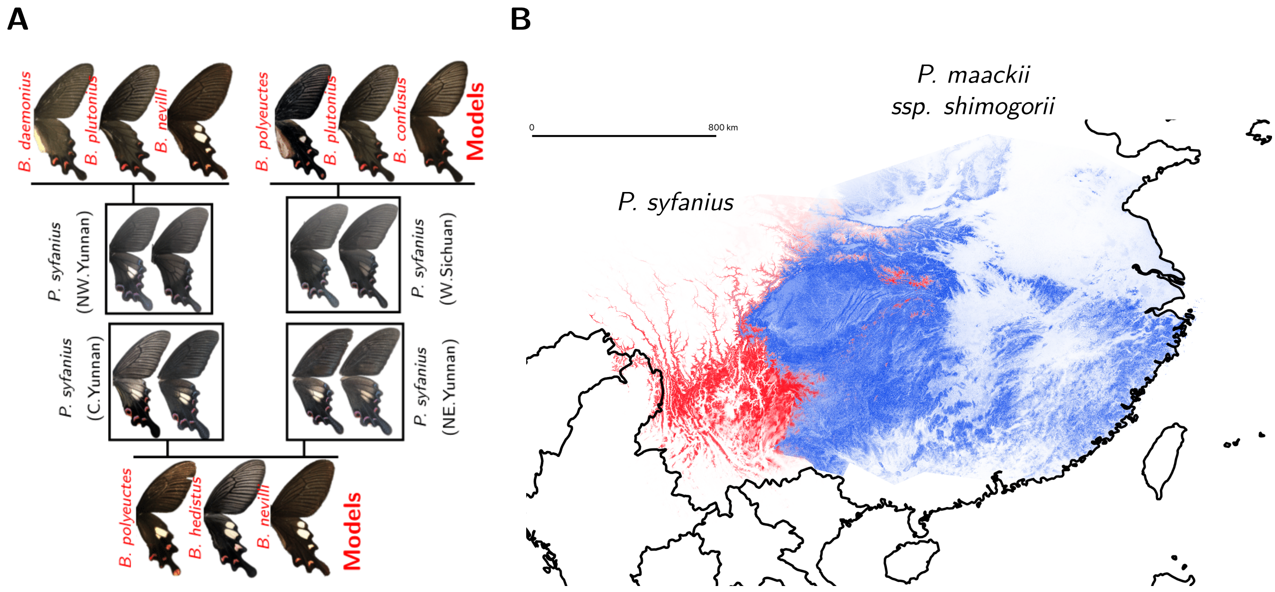

**C**

| *P. syfanius* | *P. maackii* |
|---|---|
| *Zanthoxylum acanthopodium* (major)<br>*Zanthoxylum armatum* (minor) | *Phellodendron amurense* (major)<br>*Zanthoxylum ailanthoides* (major)<br>*Tetradium glabrifolium* (major)<br>*Zanthoxylum armatum* (minor) |

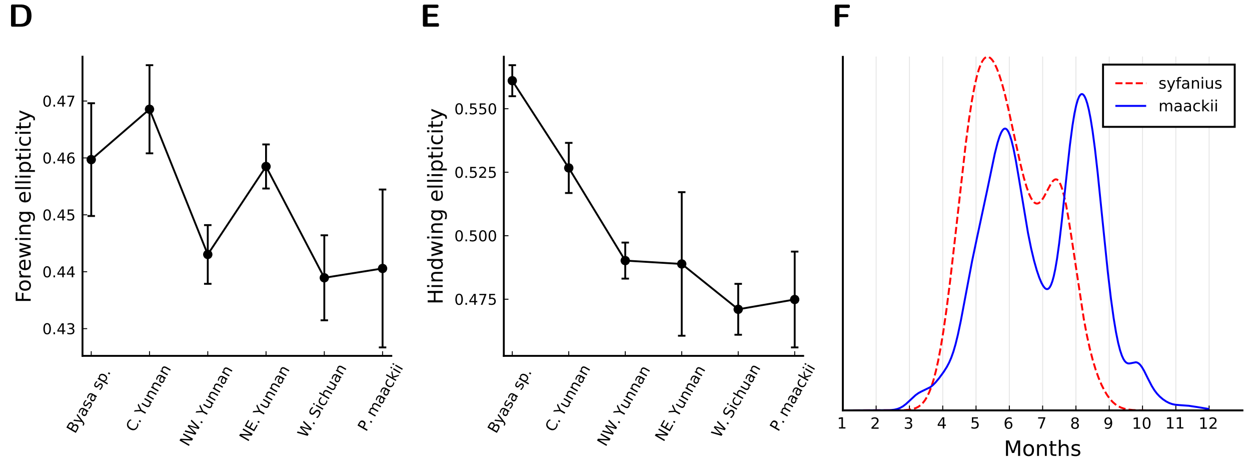

**Figure S1:** Ecology of the study system. **(A)** The mimicry polymorphism in *P. syfanius* and *P. maackii* (solid boxes). The occurrence of spotted and spotless forms matches local forms of poisonous swallowtails in the genus *Byasa*. **(B)** The geographic distribution of *P. syfanius* (red) and *P. maackii ssp. shimogorii* (blue) predicted by MaxEnt. Scale-bar represents 800km. **(C)** Host plant usage. Data are based on field observation. **(D,E)** Wing shape mimicry in *P. syfanius*. Specimens used for the study of wing shapes are from Tianzhu Xiong's personal collection and The University Museum (The University of Tokyo, Japan). Wing boundaries were extracted by a custom MATLAB script. For each wing, we fit an ellipse using the elliptic Fourier approximation for a closed boundary (Kuhl and Giardina, 1982) provided by the python package PyEFD. Ellipticity ($1 - r_{\mathrm{short}}/r_{\mathrm{long}}$) was used as a coarse measure of how wings of *P. syfanius* elongate to mimic those of the poisonous genus *Byasa*. **(F)** Seasonal distributions of adults. Records with verifiable time information (must have months, but we also kept records with missing days or years) were used. Records explicitly mentioning larvae, pupae or eggs were removed, and records from December to February were also removed since adults do not occur in winter. Together, there are 62 *P. syfanius* records and 3,138 *P. maackii* records passing all filters. Then we estimated kernel density of variable (month + day/30). When day is missing, it was randomly picked among 1 to 30. Results show the average kernel density from 1,000 repeated estimations.

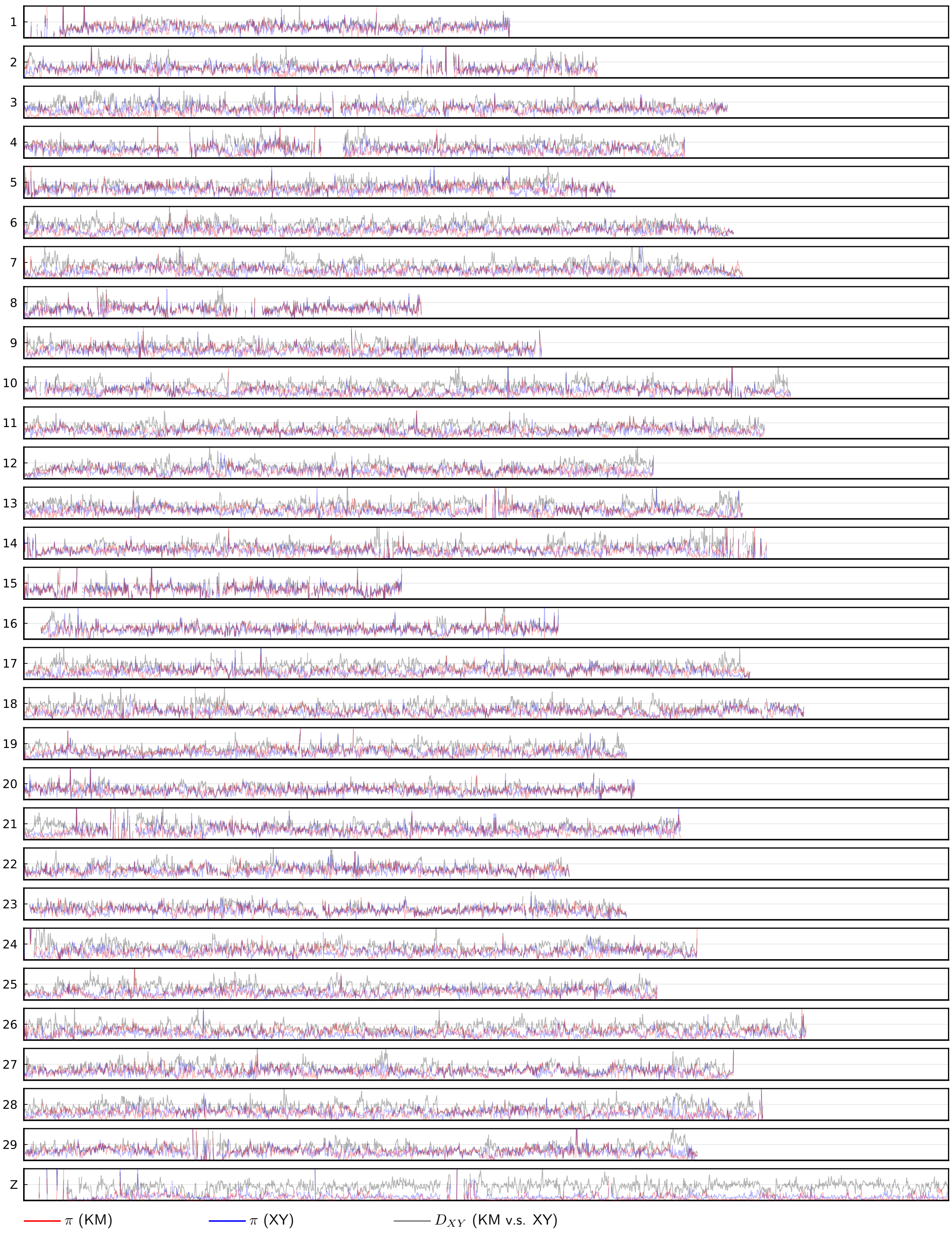

**Figure S2:** $D_{XY}$ and $\pi$ on chromosomes. This series of plots include genetic diversity $\pi$ (red: population KM; blue: population XY) and absolute sequence divergence $D_{XY}$ (gray: between populations XY and KM). $D_{XY}$ and $\pi$ are calculated on 10kb non-overlapping windows. The horizontal axis has range (0bp,$18 \times 10^6$bp), and the vertial axis has range (0,0.03). Empty regions are masked due to abnormal coverage.

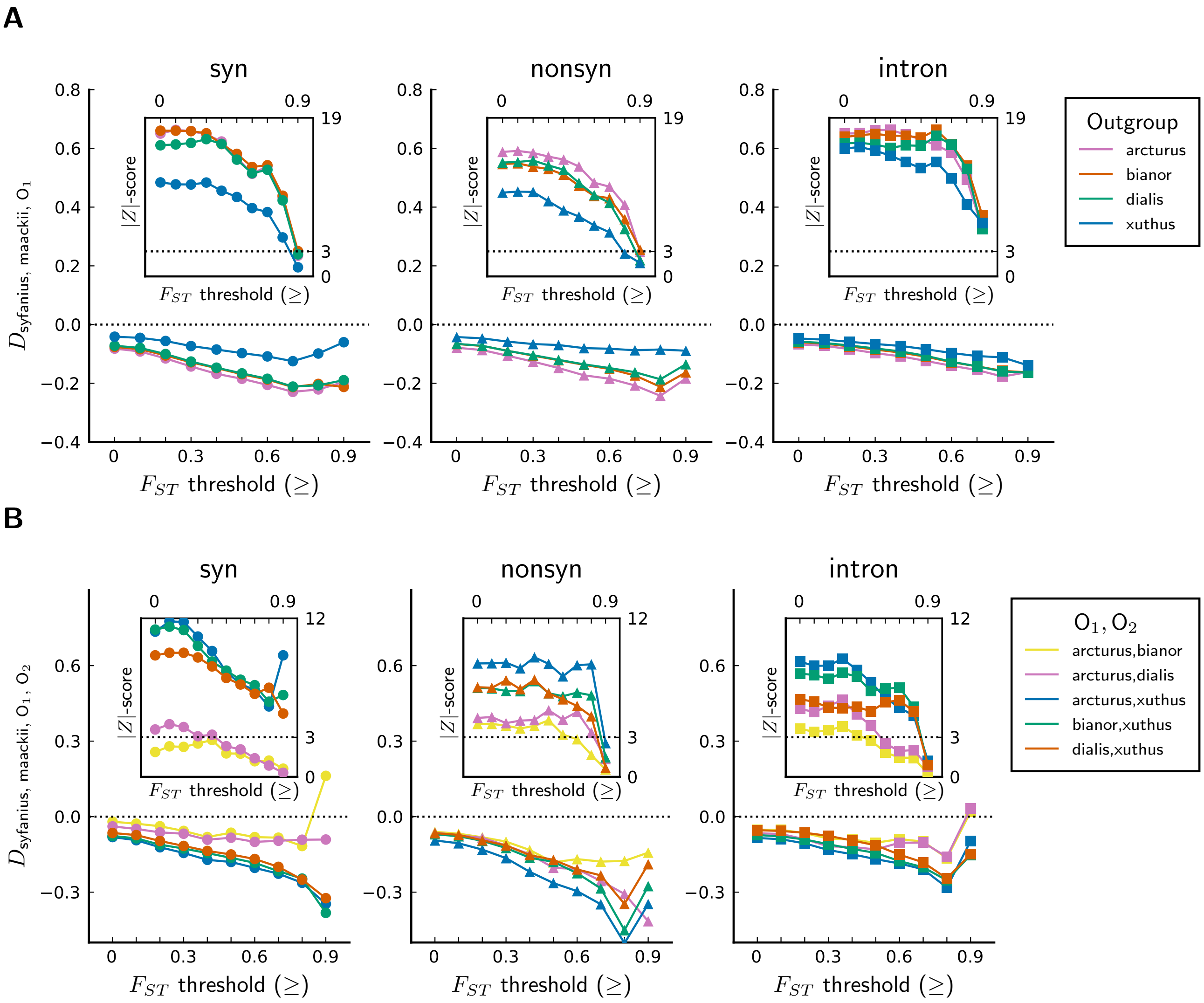

**Figure S3:** $D_3$, $D_4$ statistics and their $Z$-scores.

**Table S2:** Copy number abnormality in post-filtered SNPs data. When the average coverage of a coding sequence (CDS) exceeds twice of the median coverage of all CDSs in the genome, there is evidence for increased copy number (ICN) of that gene. We found that coding sequences or SNPs with ICN constitute a very small fraction of the entire data set, and will have negligible effect on site patterns.

| Population | #CDS with ICN unique to the population | #CDS with ICN shared between the populations |
|---|---|---|
| XY (*maackii*) | 38 | 92 |
| KM (*syfanius*) | 105 | |
| Number of SNPs with ICN from all coding sequences | 1993 | |
| Number of total SNPs from all coding sequences | 1060525 | |

**Table S3:** Results of the Wilcoxon rank-sum test. For each pair of individuals $(i, j)$, where $i$ is *P. maackii* and $j$ is *P. syfanius*, the test compares if branches leading to $i$ ($L_i$) is significantly longer than branches leading to $j$ ($L_j$). $L_i$ and $L_j$ are paired data (4776 measurements in total). $p$-values are single-sided such that rejecting the null hypothesis ($\langle L_i - L_j \rangle = 0$) suggests $\langle L_i - L_j \rangle \gg 0$. Branch lengths are defined as the distances on inferred trees from each tip to the most-recent common ancestor of all *P. syfanius*+*P. maackii* individuals.

| Individuals | Number of trees | Statistic | $p$-value |
|---|---|---|---|
| 10,4 | 4776 | 7703327.0 | $4.685783804277146\times10^{-98}$ |
| 10,5 | 4776 | 7711927.0 | $7.004020220058466\times10^{-99}$ |
| 10,8 | 4776 | 7786921.0 | $3.107578588938438\times10^{-106}$ |
| 10,2 | 4776 | 8466401.0 | $4.343367490357223\times10^{-185}$ |
| 10,3 | 4776 | 7998577.0 | $1.960298558161251\times10^{-128}$ |
| 10,7 | 4776 | 8097279.0 | $1.6161364665311736\times10^{-139}$ |
| 10,9 | 4776 | 8040167.0 | $4.784474051617971\times10^{-133}$ |
| 1,4 | 4776 | 8107709.0 | $1.028298630798444\times10^{-140}$ |
| 1,5 | 4776 | 8125705.0 | $8.549618821648854\times10^{-143}$ |
| 1,8 | 4776 | 8192570.0 | $1.1719789238813815\times10^{-150}$ |
| 1,2 | 4776 | 8845358.0 | $1.2023319895502828\times10^{-238}$ |
| 1,3 | 4776 | 8385222.0 | $1.6558753129750565\times10^{-174}$ |
| 1,7 | 4776 | 8501824.0 | $8.27364683414231\times10^{-190}$ |
| 1,9 | 4776 | 8435804.0 | $4.587014907065287\times10^{-181}$ |
| 11,4 | 4776 | 8619969.0 | $5.779268833919884\times10^{-206}$ |
| 11,5 | 4776 | 8674639.0 | $1.1426897360762765\times10^{-213}$ |
| 11,8 | 4776 | 8695345.0 | $1.2660684490436066\times10^{-216}$ |
| 11,2 | 4776 | 9292816.0 | $1.0203101578327\times10^{-310}$ |
| 11,3 | 4776 | 8906422.5 | $6.44032512136094\times10^{-248}$ |
| 11,7 | 4776 | 9011758.0 | $2.469476039690487\times10^{-264}$ |
| 11,9 | 4776 | 8928703.0 | $2.4024620857572672\times10^{-251}$ |
| 6,4 | 4776 | 8691441.0 | $4.580452175666067\times10^{-216}$ |
| 6,5 | 4776 | 8690225.0 | $6.845989175075731\times10^{-216}$ |
| 6,8 | 4776 | 8750724.0 | $1.2544536419335296\times10^{-224}$ |
| 6,2 | 4776 | 9326087.0 | 0.0 |
| 6,3 | 4776 | 8968414.0 | $1.6234720872664837\times10^{-257}$ |
| 6,7 | 4776 | 9013391.5 | $1.3611512438572988\times10^{-264}$ |
| 6,9 | 4776 | 9007066.0 | $1.3767244403926598\times10^{-263}$ |

| Individual index | Sample name |
|---|---|
| 1 | TXSC20180711006 |
| 2 | TXSC20180714006 |
| 3 | TXSC20180714008 |
| 4 | TXSC20180718016 |
| 5 | TXSC20180718009 |
| 6 | TXSC20180713001 |
| 7 | TXSC20180713007 |
| 8 | TXSC20180718001 |
| 9 | TXSC20180713008 |
| 10 | TXSC20180727005 |
| 11 | TXSC20180711011 |

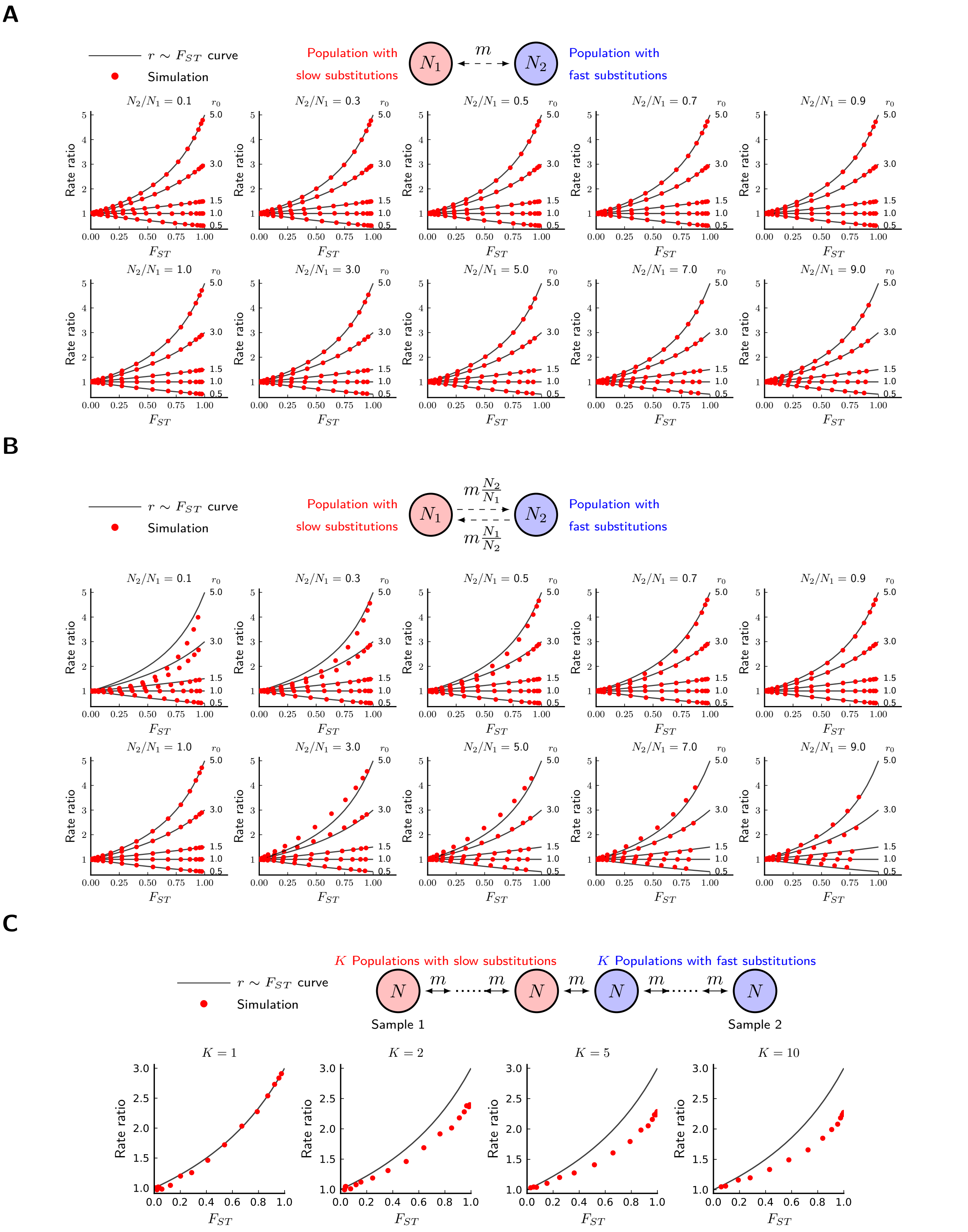

**Figure S4:** Results from equilibrium coalescent models. Fixed parameters are: $N_1 = N = 10^4/6$, $\mu_1 = 3 \times 10^{-5}$ (substitution rate in the red species), $m = (0.1)^{\mathrm{Array}(1.4:0.25:5.4)}$. **(A)** Symmetric migration models. The curve appears to be exact. ($10^6$ repetitions). **(B)** Conservative migration models. The curve is still robust, although less accurate. ($10^6$ repetitions). **(C)** Stepping-stone models. Within-species population structure dampens observed relationship below the theoretical curve. ($10^4$ repetitions).

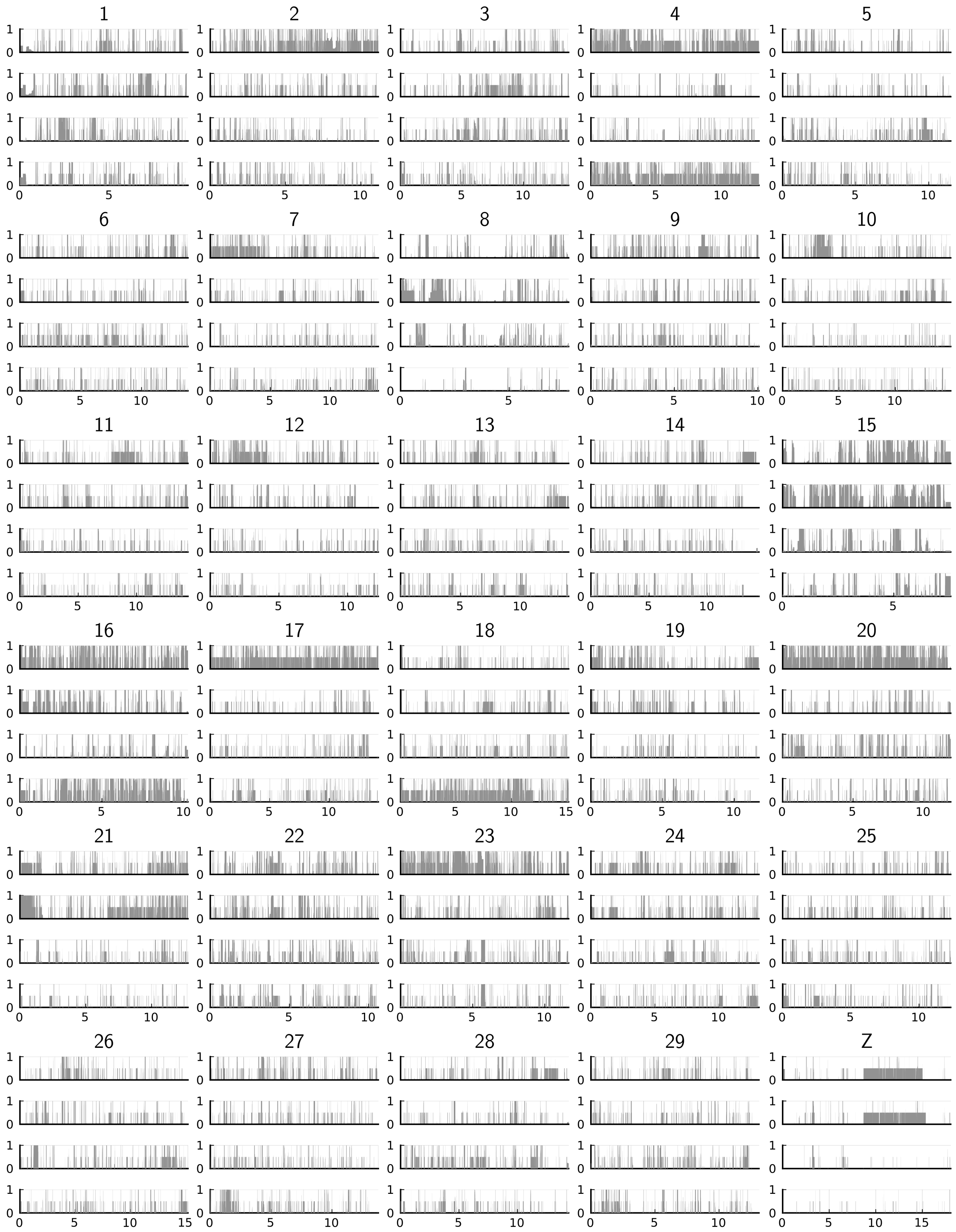

**Figure S5:** A single run of the local ancestry estimation software ELAI on chromosomes 1-30 (generation parameter: 5000) of four diploid individuals from the hybrid population WN (see Table. S1 for sample names). Gray represents the ancestry contribution from *P. maackii*.

### 1 Measuring ancestry randomness on genomic windows with a small sample size

#### 1.1 Representation of ancestry on a hybrid chromosome

The ancestry of a hybrid depends on the pure reference populations. The following assumptions are used throughout this section:

- There is a finite number of pure reference populations indexed by $k \in \{1, 2, \cdots, K\}$. In most cases, we are only interested in $K = 2$, for instance, a hybrid zone between two lineages.
- The chromosome has so many sites so that it can be treated as a contiguous rod with length $L$. $l \in [0, L]$ is the index of positions on a chromosome.
- The species' ploidy is $n_p$ (completely phased data $\Leftrightarrow n_p = 1$). When not specified, we assume the data is unphased, so that the ancestry on a particular chromosome always refers to the ancestry on a collection of $n_p$ homologous chromosomes.
- Ancestry can be inferred. In practice, regions with low density of informative SNPs will make inference difficult.

As "ancestry" is a categorical variable (i.e., there is no intrinsic order among the reference populations), to quantify the correlation of ancestry along a chromosome, we need to map each ancestry category to some numeric values that can be used to calculate correlation.

The first choice is to use the probability of each category. For unphased data, let $p_k(l)$ be the fraction of the $n_p$-ploid chromosome at position $l$ coming from reference population $k$. The vector-valued function $\mathbf{p}(l) = (p_1(l), p_2(l), \cdots, p_K(l))^\top$ completely describes the ancestry on a single chromosome. By definition, $\sum_k p_k(l) \equiv 1$. This is the conservation of total ancestry at any position. However, by this representation, the correlation $\mathbf{p}^\top(l_1)\mathbf{p}(l_2)$, which is the product of ancestries at different positions, will have units of the square of probabilities. If a locus is to be compared with itself, then the correlation will be

$$\mathbf{p}^\top(l)\mathbf{p}(l) = \sum_k p_k^2(l) \leq 1 \tag{S1}$$

Ideally, we want the correlation of ancestry of a locus with itself to be the same regardless of its ancestry configuration. Equation (S1) does not meet this criteria as it depends on $\mathbf{p}$. Here we borrow some concepts from signal-processing theory. The conservation of total ancestry means that ancestry is more analogous to the "energy" of a signal, which is in units of [squared-signal], rather than the signal itself. Therefore, we could instead define a second type of ancestry representation by taking the square roots of probabilities.

**Definition 1** (Spherical representation of ancestry)**.** . *If the probability representation of ancestry is $\mathbf{p}(l)$, then the spherical representation $\mathbf{y}(l)$ takes the square-root of each element of $\mathbf{p}(l)$:*

$$\mathbf{y}(l) = (\sqrt{p_1(l)}, \sqrt{p_2(l)}, \cdots, \sqrt{p_K(l)})^\top \tag{S2}$$

Following this representation, the autocorrelation between two spherical ancestries becomes a natural measure of the similarity between two probability distributions:

**Definition 2** (Correlation between two spherical ancestries)**.** *The autocorrelation function between $\mathbf{y}(l_1)$ and $\mathbf{y}(l_2)$ is the following quantity known as the Bhattacharyya coefficient, also known as the fidelity measure in information theory:*

$$A(l_1, l_2) = \mathbf{y}^\top(l_1)\mathbf{y}(l_2) = \sum_k \sqrt{y_k(l_1)y_k(l_2)} \in [0, 1] \tag{S3}$$

Further, self-correlation is always 1:

$$A(l,l) = \mathbf{y}^\top(l)\mathbf{y}(l) = \sum_k p_k(l) \equiv 1 \tag{S4}$$

The conservation of self-correlation will be important when we decompose ancestry into its spectral components, as it will not bias our analysis to any particular region of the chromosome. There is also a geometric meaning associated with this representation. Since $\mathbf{y}(l)$ has a $L^2$-norm of 1, each ancestry configuration corresponds to a point on the unit sphere in $\mathbb{R}^K$, and different ancestries are represented by the orientation of this vector in $\mathbb{R}^K$.

In many studies we are dealing only with two reference populations, such as hybrid zones between a pair of divergent lineages. This situation allows a more compact representation of ancestry using complex numbers:

**Definition 3** (Complex representation of ancestry)**.** *If $K = 2$, define the following complex variable $z(l)$ as the representation of ancestries:*[1]

$$z(l) = \sqrt{p_1(l)} + i\sqrt{p_2(l)} \tag{S5}$$

*where $i = \sqrt{-1}$ is the imaginary number.*

A complex-valued signal $z(l)$, like any real signal, has the definition of autocorrelation:

**Definition 4** (Correlation between two complex ancestries)**.** *The correlation between two complex ancestries $z(l_1)$ and $z(l_2)$ is the following product:*

$$\begin{aligned} A(l_1, l_2) &= z(l_1)\overline{z(l_2)} \\ &= \sqrt{p_1(l_1)p_1(l_2)} + \sqrt{p_2(l_1)p_2(l_2)} + i(\sqrt{p_2(l_1)p_1(l_2)} - \sqrt{p_1(l_1)p_2(l_2)}) \end{aligned} \tag{S6}$$

By this definition, the real part $\mathrm{Re}A(l_1, l_2)$ of the autocorrelation is just the Bhattacharyya coefficient between two ancestry configurations on $l_1$ and $l_2$, which is a measure of similarity. The absolute value of the imaginary part of the autocorrelation $|\mathrm{Im}A(l_1, l_2)|$ measures the volume of the parallelogram spanned by vectors $\mathbf{y}(l_1)$ and $\mathbf{y}(l_2)$, thus it is a measure of dissimilarity. Further, the self-correlation $A(l,l)$ is also conserved for all $l$, as $\mathrm{Im}A(l,l) \equiv 0$ and $\mathrm{Re}A(l,l) \equiv 1$.

The complex representation possesses some useful properties not shared with the spherical representation, but it also limits our analysis to cases with only two reference populations. This will not pose a problem for the study of hybrid zones between a pair of parapatric lineages.

#### 1.2 Mean autocorrelation vs. hybrid index

The hybrid index is defined as the average ancestry in a hybrid from a given reference population over a set of loci (Anderson, 1949). It is usually calculated for each individual or for each chromosome. In our notation:

**Definition 5** (Hybrid index)**.** *The hybrid index over a given genomic interval $[0, L]$ for reference population $k$ is*

$$h_k = \frac{1}{L}\int_0^L p_k(l)\,\mathrm{d}l \tag{S7}$$

In contrast, the average of the spherical or the complex representation provides information about autocorrelation within the chromosome instead of the mean ancestry.

**Theorem 1** (Mean autocorrelation)**.** *With the spherical or the complex representation, the squared $L^2$-norm (or the squared modulus, when a complex representation is available) of the average signal represents the mean*

[1]This seemingly artificial definition is not the first time that a complex number is used to model a physical phenomenon. In quantumn mechanics, the quantumn wave function of a particle is represented by a complex wave with the probability of occurrence measured by the square-modulus of the wave. Here, we are also using the square-modulus of $z(l)$ to represent the total ancestry of a given locus.

*autocorrelation of the ancestry on the genomic interval* $[0, L]$.

$$a := \frac{1}{L^2} \iint\limits_{[0,L]^2} A(l_1, l_2) \, \mathrm{d}l_1 \, \mathrm{d}l_2 = \left\| \frac{1}{L} \int_0^L \mathbf{y}(l) \, \mathrm{d}l \right\|^2 = \left| \frac{1}{L} \int_0^L z(l) \, \mathrm{d}l \right|^2 \tag{S8}$$

*Proof.* For a general spherical representation $\mathbf{y}(l)$, we have

$$\begin{aligned} \left\| \frac{1}{L} \int_0^L \mathbf{y}(l) \, \mathrm{d}l \right\|^2 &= \frac{1}{L^2} \sum_k \left( \int_0^L \sqrt{p_k(l)} \, \mathrm{d}l \right)^2 = \frac{1}{L^2} \sum_k \int_0^L \sqrt{p_k(l_1)} \, \mathrm{d}l_1 \int_0^L \sqrt{p_k(l_2)} \, \mathrm{d}l_2 \\ &= \frac{1}{L^2} \iint\limits_{[0,1]^2} \sum_k \sqrt{p_k(l_1) p_k(l_2)} \, \mathrm{d}l_1 \, \mathrm{d}l_2 = \frac{1}{L^2} \iint\limits_{[0,1]^2} A(l_1, l_2) \, \mathrm{d}l_1 \, \mathrm{d}l_2 \end{aligned} \tag{S9}$$

For a complex representation $z(l)$, notice that $A(l_1, l_2) = \overline{A(l_2, l_1)}$, so

$$\frac{1}{L^2} \iint\limits_{[0,L]^2} A(l_1, l_2) \, \mathrm{d}l_1 \, \mathrm{d}l_2 = \frac{1}{L^2} \iint\limits_{[0,L]^2} \mathrm{Re} A(l_1, l_2) \, \mathrm{d}l_1 \, \mathrm{d}l_2 = \frac{1}{L^2} \iint\limits_{[0,1]^2} \sum_{k=1,2} \sqrt{p_k(l_1) p_k(l_2)} \, \mathrm{d}l_1 \, \mathrm{d}l_2 \tag{S10}$$

This guarantees that the mean autocorrelation when using a complex representation is the same as the mean autocorrelation when using a spherical representation with $K = 2$. Additionally,

$$\left| \frac{1}{L} \int_0^L z(l) \, \mathrm{d}l \right|^2 = \frac{1}{L^2} \int_0^L z(l_1) \, \mathrm{d}l_1 \overline{\int_0^L z(l_2) \, \mathrm{d}l_2} = \frac{1}{L^2} \iint\limits_{[0,L]^2} z(l_1) \overline{z(l_2)} \, \mathrm{d}l_1 \, \mathrm{d}l_2 = \frac{1}{L^2} \iint\limits_{[0,L]^2} A(l_1, l_2) \, \mathrm{d}l_1 \, \mathrm{d}l_2 \tag{S11}$$

This completes the proof. □

The quantity $a$ is a measure of the average similarity of the ancestry configurations. It does not consider the genomic position of different ancestry configurations, so it does not contain information about whether similar ancestry configurations are clustered together.

#### 1.3 The entropy of ancestry

To characterize the scale of correlation, the distance between loci is important because correlation often drops while distance between loci increases. The information about the spatial scale of correlation is retained when the full spectrum of correlation is considered within a single individual.

A second source of correlation arises when we consider the relationship between individuals. To convey the main idea, let's compare between a set of ancestry signals from an inversion and a set of ancestry signals from a regular region subject to normal recombination. For the inversion, since recombination between chromosomes of different ancestry is often completely suppressed, the ancestry signal along the inversion will be close to constant. When two haploid individuals are compared at the inverted region, they are either different everywhere in terms of ancestry, or completely the same. Comparing between diploid individuals is similar, although the difference is more finely grained due to the presence of the heterozygotes. However, in a collinear region subject to a regular rate of recombination, any ancestry signal will switch randomly between states. In the latter case, the similarity between two ancestry signals will be similar across multiple pairwise comparisons. It will be helpful to think in the principal component (PC) space spanned by all individuals' ancestry signals. Ancestry signals from an inversion will form several tight clusters in the PC space, while those from a regular region will form a single cloud with a larger dispersion.

Both sources of correlation, and hence both types of randomness, can be measured using entropy in information theory.

**Definition 6** (Shannon entropy)**.** *The Shannon entropy for a discrete probability distribution* $\{p_j\}$*,* $j \in \mathbb{Z}$*, is* *defined as*

$$S(\{p_j\}) = -\sum_{j\in\mathbb{Z}} p_j \ln p_j \tag{S12}$$

*For a continuous distribution with probability density function $p(x)$, $x \in \mathbb{R}$, the Shannon differential entropy is defined as*

$$S(p(x)) = -\int_{\mathbb{R}} p(x) \ln p(x)\,\mathrm{d}x \tag{S13}$$

*For any nonnegative series $\{p_j\}$ or nonnegative function $p(x)$, suppose they converge upon summation/integration, we use the same notations $S(\{p_j\})$ and $S(p(x))$ to denote the entropy after they are appropriately normalized to probability distributions.*

Shannon entropy (or any other entropy measure) is a useful measure of the spread of a distribution over its entire configuration space. When the distribution is concentrated (low randomness, high certainty), $S$ will be low. $S = 0$ if and only if $p_j = 1$ for some $j$.

###### 1.3.1 Entropy associated with the correlation within a single individual

**Definition 7** (Fourier spectrum of the spherical representation)**.** *For the spherical representation of ancestry on genomic interval* $[0, L]$

$$\mathbf{y}(l) = (\sqrt{p_1(l)}, \sqrt{p_2(l)}, \cdots, \sqrt{p_K(l)})^\top, \tag{S14}$$

*expand each component into its Fourier series:*

$$\sqrt{p_k(l)} = \sum_{n=-\infty}^{+\infty} \hat{p}_{k,n} e^{i\frac{2\pi}{L}nl} \tag{S15}$$

*The folded Fourier spectrum is defined as*

$$\zeta_n = \begin{cases} 2\sum_{k=1}^{K} |\hat{p}_{k,n}|^2 & (n > 0) \\ \sum_{k=1}^{K} |\hat{p}_{k,0}|^2 & (n = 0) \end{cases} \tag{S16}$$

**Definition 8** (Fourier spectrum of the complex representation)**.** *For the complex representation of ancestry on genomic interval* $[0, L]$

$$z(l) = \sqrt{p_1(l)} + i\sqrt{p_2(l)}, \tag{S17}$$

*expand $z(l)$ into its Fourier series:*

$$z(l) = \sum_{n=-\infty}^{+\infty} Z_n e^{i\frac{2\pi}{L}nl} \tag{S18}$$

*The folded Fourier spectrum is defined as*

$$\zeta_n = \begin{cases} |Z_n|^2 + |Z_{-n}|^2 & (n > 0) \\ |Z_0|^2 & (n = 0) \end{cases} \tag{S19}$$

The following theorem guarantees that the folded spectrum $\zeta_n$ is the same for bi-ancestry signals using either representation.

**Theorem 2.** *When $K = 2$, the folded Fourier spectrum $\zeta_n$ is the same for $\mathbf{y}(l) = (\sqrt{p_1(l)}, \sqrt{p_2(l)})^\top$ and $z(l) = \sqrt{p_1(l)} + i\sqrt{p_2(l)}$.*

*Proof.* Expand each component into its Fourier series:

$$\sqrt{p_1(l)} = \sum_{n=-\infty}^{+\infty} \hat{p}_{1,n} e^{i\frac{2\pi}{L}nl}$$
$$\sqrt{p_2(l)} = \sum_{n=-\infty}^{+\infty} \hat{p}_{2,n} e^{i\frac{2\pi}{L}nl} \tag{S20}$$

The folded spectrum using the spherical representation is

$$\zeta_n = \begin{cases} 2|\hat{p}_{1,n}|^2 + 2|\hat{p}_{2,n}|^2 & (n > 0) \\ |\hat{p}_{1,0}|^2 + |\hat{p}_{2,0}|^2 & (n = 0) \end{cases} \tag{S21}$$

Using the linearity of Fourier expansion, we have

$$Z_n = \hat{p}_{1,n} + i\hat{p}_{2,n} \tag{S22}$$

Thus,

$$\begin{aligned} Z_n\overline{Z_n} &= (\hat{p}_{1,n} + i\hat{p}_{2,n})(\overline{\hat{p}_{1,n}} - i\overline{\hat{p}_{2,n}}) = \hat{p}_{1,n}\overline{\hat{p}_{1,n}} + \hat{p}_{2,n}\overline{\hat{p}_{2,n}} + i\hat{p}_{2,n}\overline{\hat{p}_{1,n}} - i\hat{p}_{1,n}\overline{\hat{p}_{2,n}} \\ &= \hat{p}_{1,n}\overline{\hat{p}_{1,n}} + \hat{p}_{2,n}\overline{\hat{p}_{2,n}} - 2\mathrm{Im}(\hat{p}_{2,n}\overline{\hat{p}_{1,n}}) \end{aligned} \tag{S23}$$

Since the Fourier coefficients of a real function at opposite frequencies are complex conjugates to each other, we have

$$Z_{-n}\overline{Z_{-n}} = \hat{p}_{1,-n}\overline{\hat{p}_{1,-n}} + \hat{p}_{2,-n}\overline{\hat{p}_{2,-n}} - 2\mathrm{Im}(\hat{p}_{2,-n}\overline{\hat{p}_{1,-n}}) = \overline{\hat{p}_{1,n}}\hat{p}_{1,n} + \overline{\hat{p}_{2,n}}\hat{p}_{2,n} - 2\mathrm{Im}(\overline{\hat{p}_{2,n}}\hat{p}_{1,n}) \tag{S24}$$

Finally,

$$|Z_n|^2 + |Z_{-n}|^2 = Z_n\overline{Z_n} + Z_{-n}\overline{Z_{-n}} = 2\hat{p}_{1,n}\overline{\hat{p}_{1,n}} + 2\hat{p}_{2,n}\overline{\hat{p}_{2,n}} - 2\mathrm{Im}(\hat{p}_{2,n}\overline{\hat{p}_{1,n}} + \overline{\hat{p}_{2,n}}\hat{p}_{1,n}) \tag{S25}$$

Since $\hat{p}_{2,n}\overline{\hat{p}_{1,n}} + \overline{\hat{p}_{2,n}}\hat{p}_{1,n}$ is real, the last term of the previous equation becomes zero. For the zero-th component $\zeta_0$, as both $\hat{p}_{1,0}$ and $\hat{p}_{2,0}$ are real, there is no difference between the two representations. This completes the proof. □

A nice property of the Fourier spectrum $\zeta_n$ is that it can be interpreted as a probability distribution of ancestry among different frequency components. By Parseval's theorem, it is easy to verify that $\sum_n \zeta_n = \frac{1}{L}\int_0^L (\sum_k p_k(l))\,\mathrm{d}l = 1$. From the Wiener-Khinchin theorem, the unfolded spectrum forms a Fourier transform pair with the autocorrelation function of the original signal. The significance of the Wiener-Khinchin theorem is that information about the autocorrelation of the original signal can now be extracted from the Fourier spectrum $\zeta_n$.

**Definition 9** (Within-individual entropy)**.** *Let $S_w = -\sum_n \zeta_n \ln \zeta_n$. $S_w$ is the Shannon entropy of the folded Fourier spectrum $\zeta_n$. As $S_w$ captures the correlation structure within each individual, we also call it the within-individual entropy.*

Formally, we have the following uncertainty principle relating the within-individual entropy to the scale of autocorrelation.

**Theorem 3** (Entropic uncertainty)**.** *Let $A(l) = \frac{1}{L}\int_0^L \mathbf{y}^\top(x)\mathbf{y}(x+l)\,\mathrm{d}x$ be the average autocorrelation at scale $l$ ($0 \le l \le L$) for the spherical ancestry. Here, $\mathbf{y}$ is understood as a periodic function of period $L$. The following inequality holds:*

$$S_w = S(\{\zeta_n\}) \ge \int_0^L \frac{A^2(l)}{Q} \ln \frac{A^2(l)}{Q}\,\mathrm{d}l + \left(\frac{1}{L}\int_0^L A(l)\,\mathrm{d}l - 1\right) \ln 2 + \ln L \tag{S26}$$

*where $Q = \int_0^L A^2(l)\,\mathrm{d}l$ is a normalization factor. Note that the right-hand-side of the inequality is invariant under linearly re-scaling $A(l)$ to a different interval $[0, L']$. Thus, we can also write the inequality compactly, supposing $A(l)$ has been rescaled to $[0,2]$, as*

$$S_w \geq -S(A^2(l)) + a\ln 2, \tag{S27}$$

*where $a$ is the average autocorrelation defined in Eq. S8.*

*Proof.* (i) In this part, we establish the Wiener-Khinchin relation that $A(l)$ expands into a Fourier series with coefficients $\eta_n = \sum_{k=1}^K |\hat{p}_{k,n}|^2$:

$$A(l) = \sum_{n=-\infty}^{+\infty}\sum_{k=1}^{K} |\hat{p}_{k,n}|^2 e^{i\frac{2\pi}{L}nl} \tag{S28}$$

The derivation is as follows:

$$\begin{aligned} A(l) &= \frac{1}{L}\int_0^L \sum_{k=1}^K y_k(x)y_k(x+l)\,\mathrm{d}x = \sum_{k=1}^K \frac{1}{L}\int_0^L \left(\sum_{n=-\infty}^{+\infty} \hat{p}_{k,n}e^{i\frac{2\pi}{L}nx}\right)\left(\sum_{n=-\infty}^{+\infty} \hat{p}_{k,n}e^{i\frac{2\pi}{L}n(x+l)}\right)\mathrm{d}x \\ &= \sum_{k=1}^K\sum_{n=-\infty}^{+\infty} e^{i\frac{2\pi}{L}nl}\frac{1}{L}\int_0^L \hat{p}_{k,n}\hat{p}_{k,-n}\,\mathrm{d}x = \sum_{n=-\infty}^{+\infty}\sum_{k=1}^K |\hat{p}_{k,n}|^2 e^{i\frac{2\pi}{L}nl} = \sum_{n=-\infty}^{+\infty}\eta_n e^{i\frac{2\pi}{L}nl} \end{aligned} \tag{S29}$$

(ii) Let $\int_0^L A^2(l)\,\mathrm{d}l = Q$. It is obvious that $\{\eta_n/\sqrt{Q}\}$ are Fourier series coefficients of $A(l)/\sqrt{Q}$, they obey the Hausdorff-Young inequality

$$\left(\sum_{n=-\infty}^{+\infty}\left|\eta_n/\sqrt{Q}\right|^{q'}\right)^{\frac{1}{q'}} \leq \left(\int_0^1 \left|A(xL)/\sqrt{Q/L}\right|^q \mathrm{d}x\right)^{\frac{1}{q}}, \tag{S30}$$

where $q \in (1,2]$ and $1/q' + 1/q = 1$. Let $\phi(q) = \left(\int_0^1 \left|A(xL)/\sqrt{Q/L}\right|^q \mathrm{d}x\right)^{\frac{1}{q}} - \left(\sum_{n=-\infty}^{+\infty}\left|\eta_n/\sqrt{Q}\right|^{q'}\right)^{\frac{1}{q'}}$. Since Fourier series preserve the 2-norm, $\phi(2) = 0$, and since $\phi(q) \geq 0$ for $q \in (1,2]$, we have $\phi'(2) \leq 0$. This translates into

$$\begin{aligned} &\left(\int_0^1 \left|A(xL)/\sqrt{Q/L}\right|^2 \mathrm{d}x\right)^{\frac{1}{2}} \left\{-\frac{1}{4}\ln\int_0^1 \left|A(xL)/\sqrt{Q/L}\right|^2 \mathrm{d}x + \frac{1}{2}\frac{\int_0^1 \left|A(xL)/\sqrt{Q/L}\right|^2 \ln\left|A(xL)/\sqrt{Q/L}\right| \mathrm{d}x}{\int_0^1 \left|A(xL)/\sqrt{Q/L}\right|^2 \mathrm{d}x}\right\} \\ &- \left(\sum_{n=-\infty}^{+\infty}\left|\eta_n/\sqrt{Q}\right|^2\right)^{\frac{1}{2}} \left\{\frac{1}{4}\ln\sum_{-\infty}^{+\infty}\left|\eta_n/\sqrt{Q}\right|^2 - \frac{1}{2}\frac{\sum_{n=-\infty}^{+\infty}\left|\eta_n/\sqrt{Q}\right|^2\ln\left|\eta_n/\sqrt{Q}\right|}{\sum_{n=-\infty}^{+\infty}\left|\eta_n/\sqrt{Q}\right|^2}\right\} \leq 0, \end{aligned} \tag{S31}$$

which yields

$$S(\{\eta_n^2\}) = -\sum_{n=-\infty}^{+\infty}\frac{\eta_n^2}{Q}\ln\frac{\eta_n^2}{Q} \geq \int_0^L \frac{A^2(l)}{Q}\ln\frac{A^2(l)}{Q}\,\mathrm{d}l + \ln L \tag{S32}$$

Note that the quantity $\int_0^L \frac{A^2(l)}{Q}\ln\frac{A^2(l)}{Q}\,\mathrm{d}l + \ln L$ is actually independent of $L$ upon re-scaling.

(iii) Next, we show that $S(\{\eta_n\}) \geq S(\{\eta_n^2\})$. Since $\sum_n \eta_n = 1$ and $\eta_n$ is nonnegative, we can always re-order them into a descending series $\hat{\eta}_n$ ($n \geq 0$) with the same entropy as $S(\{\eta_n\})$ (because entropy is permutation-invariant). The result follows as long as $S(\{\hat{\eta}_n\}) \geq S(\{\hat{\eta}_n^2\})$. Let $\beta_n = \hat{\eta}_{n+1}/\hat{\eta}_n \leq 1$. The two series, after normalization, can be written as

$$\begin{aligned} \{\hat{\eta}_n\} &= \hat{\eta}_0,\ \hat{\eta}_0\beta_0,\ \hat{\eta}_0\beta_0\beta_1,\ \hat{\eta}_0\beta_0\beta_1\beta_2,\ \cdots \\ \{\hat{\eta}_n^2/Q\} &= \frac{\hat{\eta}_0^2}{Q},\ \frac{\hat{\eta}_0^2}{Q}\beta_0^2,\ \frac{\hat{\eta}_0^2}{Q}\beta_0^2\beta_1^2,\ \frac{\hat{\eta}_0^2}{Q}\beta_0^2\beta_1^2\beta_2^2,\ \cdots \end{aligned} \tag{S33}$$

Consequently,

$$\hat{\eta}_0(1+\beta_0+\beta_0\beta_1+\cdots)=\frac{\hat{\eta}_0^2}{Q}(1+\beta_0^2+\beta_0^2\beta_1^2+\cdots)=1 \tag{S34}$$

This implies that $\hat{\eta}_0 \leq \hat{\eta}_0^2/Q$. Suppose there exists $j \geq 0$ such that

$$\hat{\eta}_0(1+\beta_0+\beta_0\beta_1+\cdots+\beta_0\beta_1\cdots\beta_j)>\frac{\hat{\eta}_0^2}{Q}(1+\beta_0^2+\beta_0^2\beta_1^2+\cdots+\beta_0^2\beta_1^2\cdots\beta_j^2) \tag{S35}$$

Then there exists $j' \leq j$ such that $\hat{\eta}_0\beta_0\beta_1\cdots\beta_{j'} > \frac{\hat{\eta}_0^2}{Q}\beta_0^2\beta_1^2\cdots\beta_{j'}^2$. So for any $j \geq j'$, we have $\hat{\eta}_0\beta_0\beta_1\cdots\beta_j > \frac{\hat{\eta}_0^2}{Q}\beta_0^2\beta_1^2\cdots\beta_j^2$. The difference

$$\Delta_j=\hat{\eta}_0(1+\beta_0+\beta_0\beta_1+\cdots+\beta_0\beta_1\cdots\beta_j)-\frac{\hat{\eta}_0^2}{Q}(1+\beta_0^2+\beta_0^2\beta_1^2+\cdots+\beta_0^2\beta_1^2\cdots\beta_j^2) \tag{S36}$$

will always be positive and monotonically increases for any $j \geq j'$. This is not consistent with the fact that $\lim_{j\to\infty}\Delta_j = 0$. Thus, we conclude that for any $j$, the partial sum follows the inequality

$$\hat{\eta}_0(1+\beta_0+\beta_0\beta_1+\cdots+\beta_0\beta_1\cdots\beta_j)\leq\frac{\hat{\eta}_0^2}{Q}(1+\beta_0^2+\beta_0^2\beta_1^2+\cdots+\beta_0^2\beta_1^2\cdots\beta_j^2) \tag{S37}$$

This is to say that infinite sequence $\{\hat{\eta}_0^2/Q\}$ majorizes $\{\hat{\eta}_n\}$. By Theorem 2.2 of (Li and Busch, 2013), $S(\{\hat{\eta}_n\}) \geq S(\{\hat{\eta}_n^2\})$, and so $S(\{\eta_n\}) \geq S(\{\eta_n^2\})$

(iv) Finally, since $\zeta_n = \eta_n + \eta_{-n} = 2\eta_n$ for $n \geq 1$, we have

$$S(\{\zeta_n\})=-\zeta_0\ln\zeta_0-\sum_{n\geq1}2\eta_n\ln(2\eta_n)=-\zeta_0\ln\zeta_0-\sum_{n\geq1}(\eta_n+\eta_{-n})(\ln\eta_n+\ln 2)=S(\{\eta_n\})-(1-\eta_0)\ln 2, \tag{S38}$$

which is equivalent to

$$S(\{\zeta_n\})+\left(1-\frac{1}{L}\int_0^L A(l)\,\mathrm{d}l\right)\ln 2=S(\{\eta_n\}) \tag{S39}$$

Combining previous four steps yields the result of the theorem.

□

###### 1.3.2 Entropy associated with the correlation between individuals

**Definition 10** (Entropy of a linear operator)**.** *Let $\mathcal{L}$ be a compact self-adjoint linear operator on a Hilbert space $\mathcal{H}$. If $\mathcal{L}$ has a countable set of eigenvalues $\{\nu_i\}$, and is positive semidefinite, then define the entropy of $\mathcal{L}$ as*

$$S(\mathcal{L}):=-\sum_i\frac{\nu_i}{\mathrm{Tr}\mathcal{L}}\ln\frac{\nu_i}{\mathrm{Tr}\mathcal{L}}, \tag{S40}$$

*where $\mathrm{Tr}\mathcal{L}=\sum_i\nu_i$ is the trace of the linear operator $\mathcal{L}$*

**Definition 11** (Mercer's spectrum)**.** *Let $A_j(l_1,l_2)$ be the autocorrelation function for individual $j$ $(1\leq j\leq J)$, and define the average autocorrelation function as*

$$\langle A\rangle(l_1,l_2)=\frac{1}{J}\sum_j A_j(l_1,l_2). \tag{S41}$$

*Since $A_j$ is Hermitian, the average $\langle A\rangle$ is also Hermitian, thus the integral operator $I_{\langle A\rangle}$ defined by $I_{\langle A\rangle}\phi(l)=\int_0^L\langle A\rangle(l,s)\phi(s)\,\mathrm{d}s$ has a series of real eigenvalues $\{\nu_j\}$ satisfying $\sum_j\nu_j=L$. $\nu_j$ is the solution to the eigenvalue problem:*

$$\int_0^L\langle A\rangle(l,s)\phi_j(s)\,\mathrm{d}s=\nu_j\phi_j(l) \tag{S42}$$

*The spectrum defined by $\{\nu_j/L\}$ is the Mercer's spectrum.*

**Definition 12** (Cross-correlation spectrum)**.** *Let the cross-correlation matrix be* $\mathbf{C} = \{c_{j,j'}\}_{J\times J}$*, where* $c_{j,j'}$ *is the average cross-correlation between individual* $j$ *and* $j'$*:*

$$c_{j,j'} = \begin{cases} \frac{1}{L}\int_0^L \mathbf{y}_j^\top(l)\mathbf{y}_{j'}(l)\,\mathrm{d}l & (\textit{for the spherical representation}) \\ \frac{1}{L}\int_0^L z_j(l)\overline{z_{j'}(l)}\,\mathrm{d}l & (\textit{for the complex representation}) \end{cases} \tag{S43}$$

*As* $\mathbf{C}$ *is Hermitian, it has a series of real eigenvalues* $\lambda_j$ *satisfying* $\sum_j \lambda_j = J$*. Let the normalized spectrum of* $\mathbf{C}$ *be* $\{\lambda_j/J\}$*, then* $\{\lambda_j/J\}$ *is defined as the cross-correlation spectrum.*

The following theorem states that when the complex representation is adopted for bi-ancestry signals, the Mercer's spectrum coincides with the cross-correlation spectrum, so that even if the Mercer's spectrum is calculated using correlation *within* individuals, both captures the correlation *between* individuals.

**Theorem 4.** *If* $\{z_j(l)\}_{j=1}^J$ *is a collection of complex bi-ancestry signals, then its Mercers' spectrum is the same as its cross-correlation spectrum.*

*Proof.* The average autocorrelation $\langle A\rangle$ is computed as

$$\langle A\rangle(l_1,l_2) = \frac{1}{J}\sum_j z_j(l_1)\overline{z_j(l_2)} \tag{S44}$$

So the integral equation (S42) becomes

$$\frac{1}{J}\sum_j z_j(l)\int_0^L \overline{z_j(s)}\phi(s)\,\mathrm{d}s = \nu\phi(l), \tag{S45}$$

where $(\nu,\phi)$ forms the solution to the above equation. Rearranging the terms, we write

$$\phi(l) = \sum_j \left[\frac{\nu^{-1}}{J}\int_0^L \overline{z_j(s)}\phi(s)\,\mathrm{d}s\right] z_j(l) = \sum_j \alpha_j z_j(l), \tag{S46}$$

where $\alpha_j$ stands for the constant inside the bracket. Substituting into the original integral equation, we have

$$\frac{1}{J}\sum_j z_j(l)\int_0^L \overline{z_j(s)}\sum_{j'}\alpha_{j'}z_{j'}(s)\,\mathrm{d}s = \nu\sum_j \alpha_j z_j(l), \tag{S47}$$

which is equivalent to

$$\sum_j\left[\sum_{j'}\alpha_{j'}\int_0^L z_{j'}(s)\overline{z_j(s)}\,\mathrm{d}s\right] z_j(l) = J\nu\sum_j \alpha_j z_j(l) \tag{S48}$$

Notice that the integral in the bracket contains the cross-correlation between $j'$ and $j$, so if the following relationship holds

$$\sum_{j'}\alpha_{j'}\frac{1}{L}\int_0^L z_{j'}(s)\overline{z_j(s)}\,\mathrm{d}s = \sum_{j'}\alpha_{j'}\overline{c_{j,j'}} = \frac{J\nu}{L}\alpha_j \tag{S49}$$

then the system has a solution of $\nu$. The above equation is equivalent to

$$\overline{\mathbf{C}}\boldsymbol{\alpha} = \frac{J\nu}{L}\boldsymbol{\alpha} \Leftrightarrow \mathbf{C}\overline{\boldsymbol{\alpha}} = \frac{J\nu}{L}\overline{\boldsymbol{\alpha}} \tag{S50}$$

Which means that $J\nu/L$ is the eigenvalue of the cross-correlation matrix $\mathbf{C}$, and since $\mathbf{C}$ is a Hermitian matrix, the existence of its real eigenvalues is guaranteed. So for each $\nu_i$, we have $\lambda_j = J\nu_j/L$, which is to say

$$\frac{\lambda_j}{J} = \frac{\nu_j}{L}, \quad \text{for } \forall\ i \tag{S51}$$

□

As both spectra coincide for the complex representation, we can define the between-individual entropy using either of them.

**Definition 13** (Between-individual entropy of the complex representation)**.** *For complex bi-ancestry signals, let* $S_b := S(I_{\langle A\rangle}) = S(\mathbf{C})$*.* $S_b$ *is the Shannon entropy characterizing the between-individual correlation.*

**Corollary 1** (Maximum $S_b$)**.** *For* $J$ *diploid individuals with the complex representation of ancestry, the largest attainable* $S_b$ *is given by*

$$\begin{aligned} S_{b,max} &= -\frac{(J-1)(1-c)}{J}\ln\frac{1-c}{J} - \frac{1+(J-1)c}{J}\ln\frac{1+(J-1)c}{J}, \\ &\approx (1-c)\ln J - (1-c)\ln(1-c) - c\ln c \quad (J \gg 1) \end{aligned} \tag{S52}$$

*where* $c = (3+2\sqrt{2})/8$*.*

*Proof.* For entropy to be large, autocorrelation within each individual must be very weak. This will occur if a sufficient amount of recombination has taken place such that the ancestry across the genome is largely independent between loci. The maximum value will occur when both parental lineages contribute equally to the hybrid ancestry, so that the hybrid index $h = 0.5$. The heterozygosity $H$ is then 0.5 following a random distribution of ancestries along the two genome copies. The off-diagonal elements in the cross-correlation matrix $\mathbf{C}$ thus take the value

$$c = 2 \times \left(\frac{1}{2} + \frac{\sqrt{2}-1}{2}\frac{1}{2}\right)^2 = \frac{3+2\sqrt{2}}{8} \tag{S53}$$

For a $J \times J$ matrix whose diagonal elements are all 1, and whose off-diagonal elements are all $c$, its normalized eigenvalues are given by $\{\frac{1+(J-1)c}{J}, \frac{1-c}{J}, \cdots, \frac{1-c}{J}\}$, which leads to the above result. □

**Corollary 2** (Minimum $S_b$)**.** $S_b = 0$ *is the minimum between-individual entropy, and is attainable if recombination is completely suppressed in the genomic interval of interest.*

*Proof.* If recombination is completely suppressed along $[0, L]$, the ancestry signal is constant along the interval in any individual with any ploidy. Hence, $A_j \equiv 1$, and $\langle A\rangle \equiv 1$. The only non-zero eigenvalue associated with a constant integral kernel is $\nu = L$, so that we have $S_b = 0$ using the Mercer's spectrum. Note that the converse is not true, because the diploid ancestry signal is unaware of phase. A region completely heterozygous may in fact has a recombination break point, and the ancestry can flip phases when crossing the break point. However, this extreme situation is unlikely as it requires two separate recombination events to have occurred at precisely the same point and in opposite directions. If the hybrid zone is old, a long track of heterozygous ancestry is therefore usually good evidence for the presence of barrier loci. □

### References

E. Anderson. *Introgressive Hybridization.* John Wiley, 1949.

M. Harada, T. Teshirogi, H. Ozawa, and M. Yago. Catalogue of the Suguru Igarashi insect collection, Part I. Lepidoptera, Papilionidae. *The University Museum, The University of Tokyo, Material Reports*, 94:1–390, 2012.

F. P. Kuhl and C. R. Giardina. Elliptic Fourier features of a closed contour. *Computer Graphics and Image Processing*, 18(3):236–258, 1982.

Y. Li and P. Busch. Von neumann entropy and majorization. *Journal of Mathematical Analysis and Applications*, 408(1):384–393, 2013.

S. Lu, J. Yang, X. Dai, F. Xie, J. He, Z. Dong, J. Mao, G. Liu, Z. Chang, R. Zhao, et al. Chromosomal-level reference genome of chinese peacock butterfly (*Papilio bianor*) based on third-generation DNA sequencing and Hi-C analysis. *GigaScience*, 8(11):giz128, 2019.

The Global Biodiversity Information Facility. Occurrence download, 2021a. URL https://doi.org/10.15468/dl.ds4c5d.

The Global Biodiversity Information Facility. Occurrence download, 2021b. URL https://doi.org/10.15468/dl.v8vbzs.

M. Yago, R. Katsuyama, M. Harada, M. Teshirogi, Y. Ogawa, S. Ishizuka, and K. Omoto. Catalogue of the Keiichi Omoto butterfly collection, Part I (Papilionidae: Baroniinae and Parnassiinae). *The University Museum, The University of Tokyo, Material Reports*, 127:1–162, 2021.
